# Characterization and genomic analysis of the Lyme disease spirochete bacteriophage ϕBB-1

**DOI:** 10.1101/2024.01.08.574763

**Authors:** Dominick R. Faith, Margie Kinnersley, Diane M. Brooks, Dan Drecktrah, Laura S. Hall, Eric Luo, Andrew Santiago-Frangos, Jenny Wachter, D. Scott Samuels, Patrick R. Secor

## Abstract

Lyme disease is a tick-borne infection caused by the spirochete *Borrelia* (*Borreliella*) *burgdorferi*. *Borrelia* species have highly fragmented genomes composed of a linear chromosome and a constellation of linear and circular plasmids some of which are required throughout the enzootic cycle. Included in this plasmid repertoire by almost all Lyme disease spirochetes are the 32-kb circular plasmid cp32 prophages that are capable of lytic replication to produce infectious virions called ϕBB-1. While the *B. burgdorferi* genome contains evidence of horizontal transfer, the mechanisms of gene transfer between strains remain unclear. While we know that ϕBB-1 transduces cp32 and shuttle vector DNA during *in vitro* cultivation, the extent of ϕBB-1 DNA transfer is not clear. Herein, we use proteomics and long-read sequencing to further characterize ϕBB-1 virions. Our studies identified the cp32 *pac* region and revealed that ϕBB-1 packages linear cp32s via a headful mechanism with preferentially packaging of plasmids containing the cp32 *pac* region. Additionally, we find ϕBB-1 packages fragments of the linear chromosome and full-length plasmids including lp54, cp26, and others. Furthermore, sequencing of ϕBB-1 packaged DNA allowed us to resolve the covalently closed hairpin telomeres for the linear *B. burgdorferi* chromosome and most linear plasmids in strain CA-11.2A. Collectively, our results shed light on the biology of the ubiquitous ϕBB-1 phage and further implicates ϕBB-1 in the generalized transduction of diverse genes and the maintenance of genetic diversity in Lyme disease spirochetes.

## Introduction

The bacterium *Borrelia* (*Borreliella*) *burgdorferi* is the causative agent of Lyme disease, the most common tick-borne disease in the Northern Hemisphere [1–3]. Lyme disease spirochetes have complex and highly fragmented genomes composed of a ∼900-kb linear chromosome and up to twenty distinct and co-existing linear and circular plasmids that are similar but not identical across the genospecies [4–6].

As a vector-borne pathogen, *B. burgdorferi* relies on the differential expression of several outer surface lipoproteins to transmit from its tick vector to a vertebrate host [7]. As such, a large fraction of the *B. burgdorferi* genome encodes outer membrane lipoproteins, mostly carried on the plasmids [6, 8, 9].

In natural populations, genetic variation in outer membrane lipoprotein alleles is associated with species-level adaptations [6, 8-10] and variation in outer membrane lipoprotein alleles across the genospecies is driven primarily by horizontal gene transfer [5, 11-21]. However, the mechanism(s) by which heterologous *B. burgdorferi* strains exchange genetic material are not well defined.

Viruses that infect bacteria (phages) are key drivers of horizontal gene transfer between bacteria [22]. The genomes of nearly all sequenced Lyme disease spirochetes include the 32-kb <u>c</u>ircular <u>p</u>lasmid (cp32) prophages (**Fig 1A and B**) [4]. The cp32s carry several outer membrane lipoprotein gene families including *bdr, mlp*, and *ospE/ospF/elp* (*erps*), which are all involved in immune evasion [23–27] and exhibit sequence variation that is consistent with historical recombination amongst cp32 plasmid isoforms [21, 28, 29]. Recent work indicates that cp32 prophages are induced in the tick midgut during a bloodmeal [9, 30, 31]. When induced, cp32 prophages undergo lytic replication where they are packaged into infectious virions designated ϕBB-1 (**Fig 1C**) [32–34].

**Figure 1.**
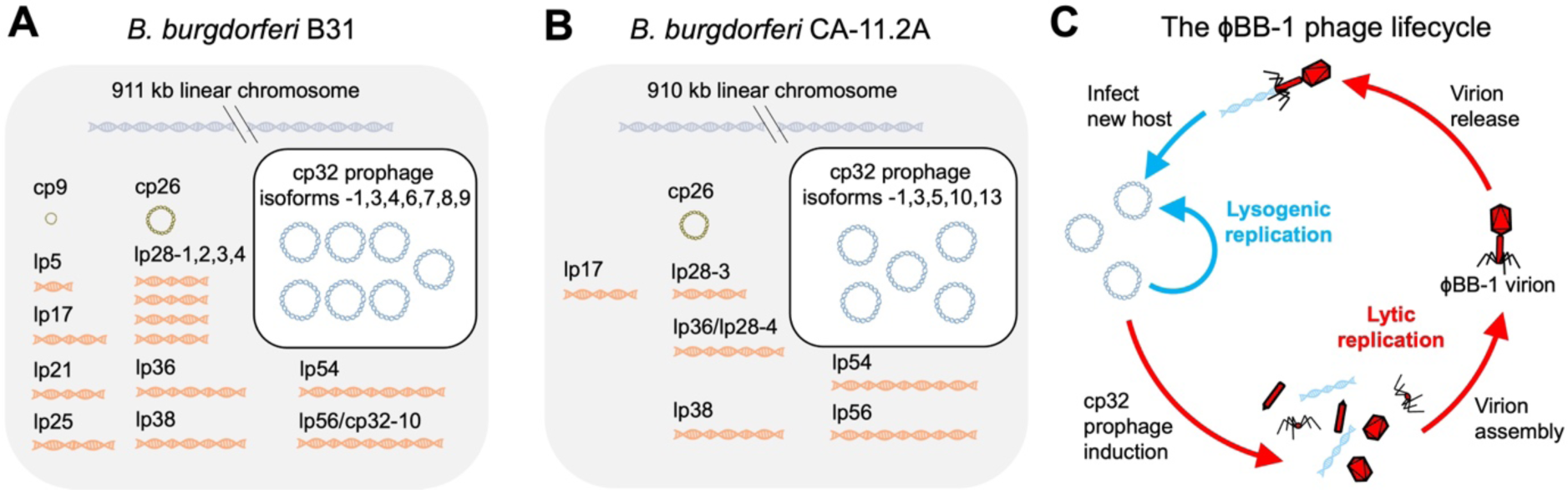
The *B. burgdorferi* genome is highly fragmented and is composed of a linear chromosome, linear and circular plasmids, and cp32 prophages. The genomes of *B. burgdorferi* strains **(A)** B31 and **(B)** CA-11.2A are shown. **(C)** The temperate ϕBB-1 phage lifecycle is depicted.

In addition to horizontally transferring phage genomes between bacterial hosts (transduction), phages frequently package and horizontally transfer pieces of the bacterial chromosome or other non-phage DNA (generalized transduction) [35]. Generalized transduction was first observed in the *Salmonella* phage P22 in the 1950s [36] and since then has been observed in numerous other phage species [35, 37-40]. ϕBB-1 is a generalized transducing phage that can horizontally transfer shuttle vectors carrying antibiotic resistance cassettes between *B. burgdorferi* strains [32]. However, to our knowledge, generalized transduction of anything other than engineered plasmids by ϕBB-1 has not been observed.

Here, we define the genetic material packaged by ϕBB-1 virions isolated from *B. burgdorferi* strain CA-11.2A. Our proteomics studies confirm that ϕBB-1 virions are composed primarily of capsid and other phage structural proteins encoded by the cp32s; however, putative phage structural proteins encoded by lp54 were also detected. Long-read sequencing reveals that ϕBB-1 virions package a variety of genetic material including cp32 isoforms that are linearized at a region immediately upstream of the *erp* locus (*ospE/ospF/elp*) and packaged into ϕBB-1 capsids via a headful genome packaging mechanism at a packaging site (*pac*). When introduced to a shuttle vector, the *pac* region promotes the packaging of shuttle vectors into ϕBB-1 virions, demonstrating the utility of ϕBB-1 as a tool to genetically manipulate Lyme disease spirochetes. Additionally, full-length contigs of cp26, lp17, lp38, lp54, and lp56 are recovered from packaged reads as are fragments of the linear chromosome. Finally, long-read sequencing of packaged DNA allowed us to fully resolve most of the covalently closed hairpin telomeres in the *B. burgdorferi* CA-11.2A genome.

Overall, this study implicates ϕBB-1 in mobilizing large portions of the *B. burgdorferi* genome, which may explain certain aspects of genome stability and diversity observed in Lyme disease spirochetes.

## Results

### ɸBB-1 phage purification, virion morphology, and proteomic analysis

In the laboratory, lytic ϕBB-1 replication (**Fig 1C**) can be induced by fermentation products such as ethanol [41, 42]. We first measured ϕBB-1 titers in early stationary-phase cultures (∼1 × 10^8^ cells/mL) of *B. burgdorferi* B31 or CA-11.2A induced with 5% ethanol, as described by Eggers *et al.* [41]. Seventy-two hours after induction, bacteria were removed by centrifugation and filtering. Virions were then purified from supernatants by chloroform extraction and precipitation with ammonium sulfate. Purified virions were treated for one hour with DNase to destroy DNA not protected within a capsid and re-chloroformed to inactivate DNAse and quantitative PCR (qPCR) was used to measure packaged cp32 copy numbers.

*B. burgdorferi* strain CA-11.2A consistently produced ∼10 times more phage than B31 (**Fig 2A**) and was selected for further study. Imaging of purified virions collected from CA-11.2A by transmission electron microscopy reveals virions with an elongated capsid and contractile tail (**Fig 2B**), which is similar to the Myoviridae morphology of ϕBB-1 virions produced by strain B31 *in vitro* [9, 43, 44] and by a human *B. burgdorferi* isolate following ciprofloxacin treatment [45].

**Figure 2.**
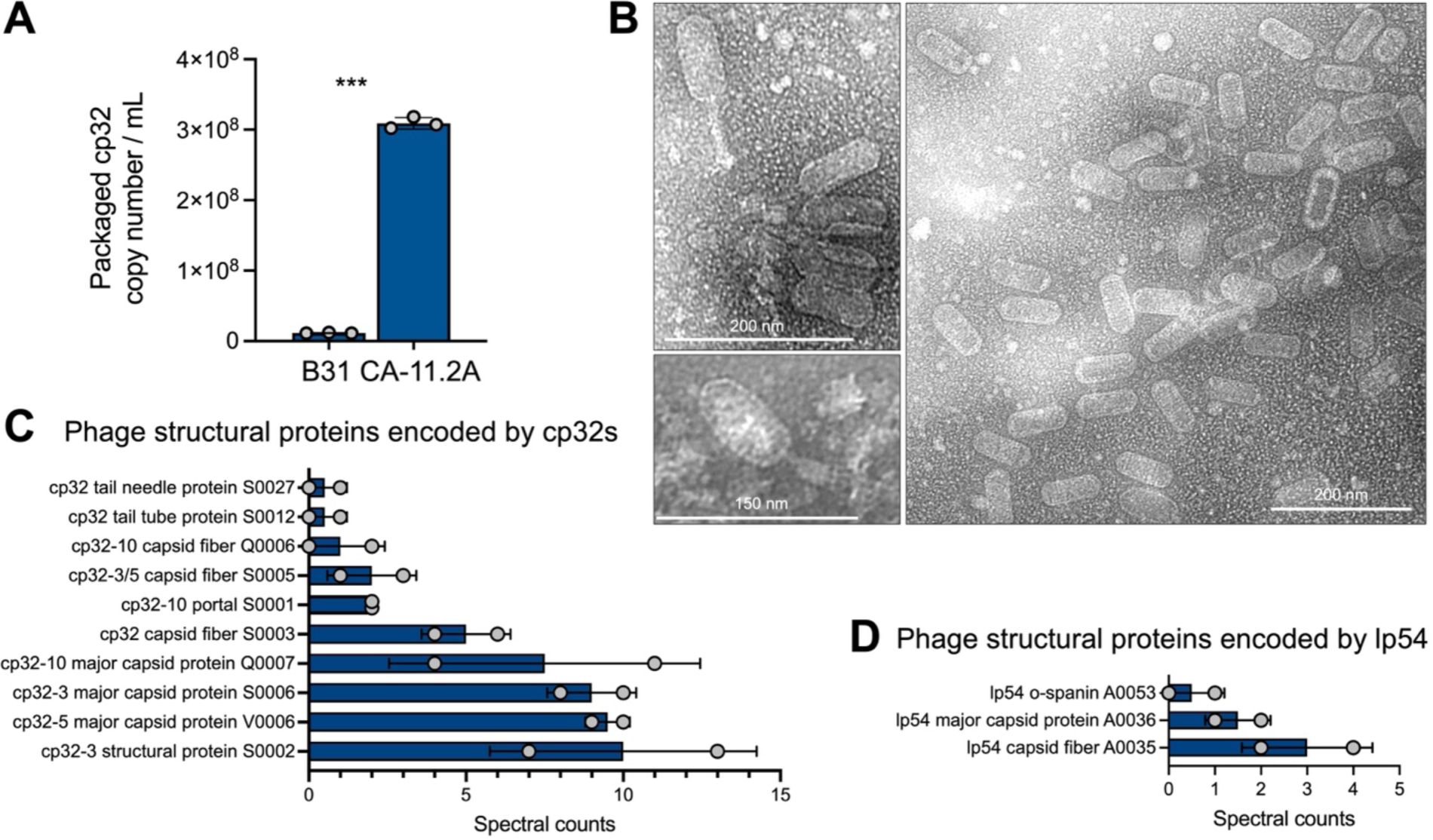
ɸBB-1 phage titer, virion morphology, and proteomic analysis. **(A)** Packaged, DNase-protected cp32 copy numbers in bacterial supernatants were measured by qPCR. Data are the SE of the mean of three experiments, \*\*\**p* <0.001. **(B)** Virions were purified from 4-L cultures of *B. burgdorferi* CA-11.2A and imaged by transmission electron microscopy. Representative images from two independent preparations are shown. **(C and D)** HPLC-MS/MS-based proteomics was used to identify proteins in two purified virion preparations. The SE of the mean of spectral counts for peptides associated with the indicated phage structural proteins are shown for each replicate. See also **Table S1** for the complete proteomics dataset.

Mass spectrometry analysis of purified virions identified ten capsid and other structural proteins encoded by the cp32s including the major capsid protein and capsid fibers (**Fig 2C, Table S1**). We also detected highly conserved predicted phage capsid proteins encoded by lp54 (**Fig 2D**). While the virions we visualized all appear to have the same elongated capsid morphology, virions with a notably smaller capsid morphology have been isolated and imaged from *B. burgdorferi* CA-11.2A [32]. These observations raise the possibility that there are multiple intact phages inhabiting the CA-11.2A genome.

### ɸBB-1 virions package portions of the *B. burgdorferi* genome

We performed long-read sequencing on DNA packaged in purified ϕBB-1 virions, as outlined in Figure 3. Although intact *B. burgdorferi* cells were removed via both centrifugation and filtration prior to chloroform treatment, there is concern that contaminating unpackaged *B. burgdorferi* chromosomal or plasmid DNA co-purifies with phage virions. To control for this, we spiked purified ϕBB-1 virions with high molecular weight (>20 kb) salmon sperm DNA (**Fig 4A**) at 1.7 µg/mL, a concentration that approximates the amount of DNA released by 3 × 10^8^ lysed bacterial cells into one milliliter of media [46]. Samples were then treated with DNase overnight followed by phage DNA extraction using a proteinase K/SDS/phenol-chloroform DNA extraction protocol [33]. Purified DNA was directly sequenced using the Nanopore MinION (long read) platform.

**Figure 3.**
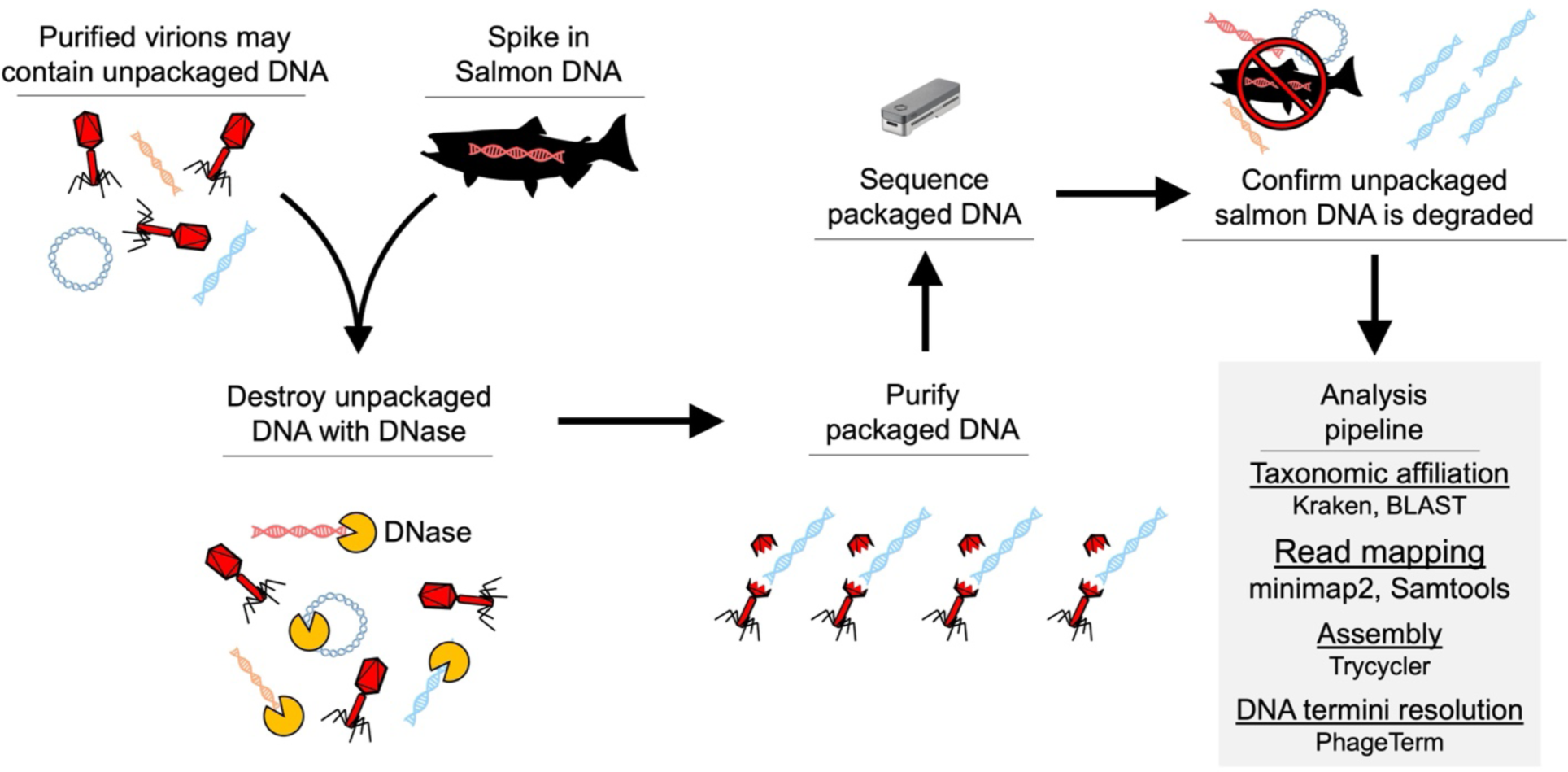
Workflow for sequencing packaged ɸBB-1 DNA.

**Figure 4.**
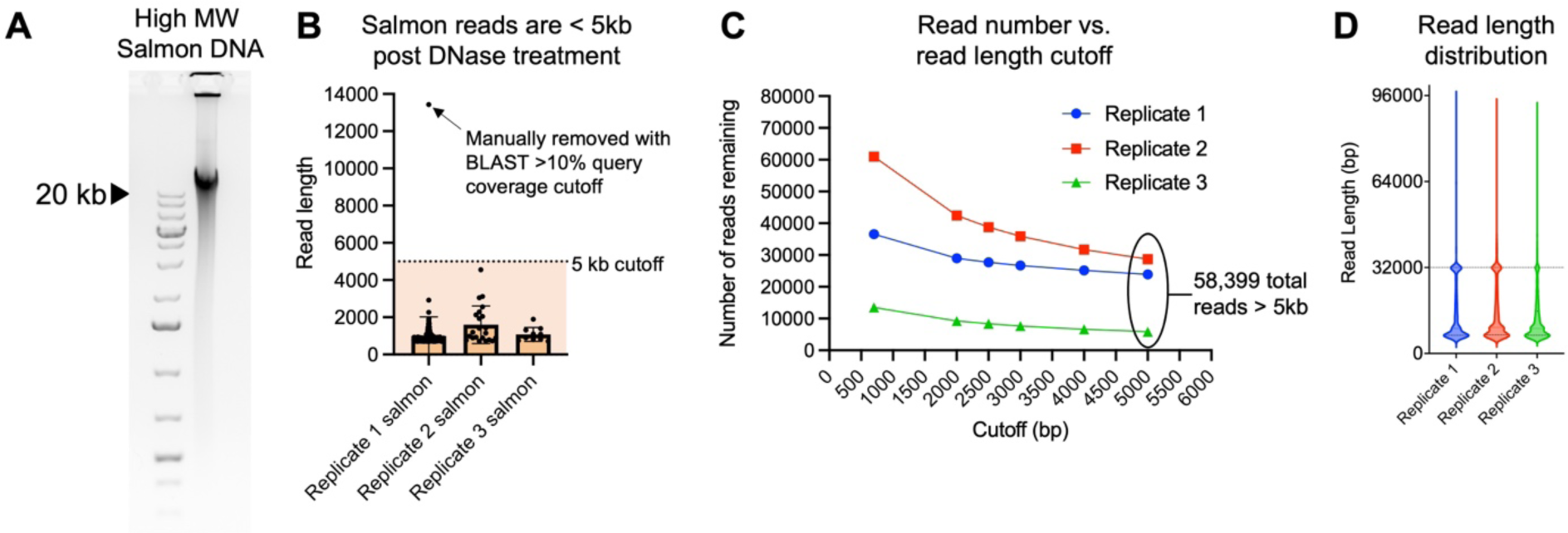
Establishing a 5kb read length cutoff to exclude unpackaged reads. **(A)** The salmon sperm DNA used to spike purified phages prior to DNase treatment was run on an agarose gel to estimate its size. Note that the majority of salmon DNA is larger than the 20-kb high molecular weight marker in the left lane. **(B)** 0.14% of 110,986 reads > 700 bp, 0.14% were classified as matching salmon sequences. Reads classified as salmon were plotted as a function of their length for each replicate. Error bars represent the SE of the mean of three replicate experiments. All reads except one (arrow) were below 5 kb in length (dashed line) with an average length of 1.2 kb. **(C)** Read length cutoff was plotted as a function of the number of reads remaining in each replicate dataset. In total, 58,399 reads remain after establishing a 5-kb cutoff. **(D)** Read length for all reads >5 kb in each replicate was plotted.

Across three replicates, we recovered a total of 110,986 nanopore reads >700 bp in length that met a minimum q-score threshold of 7. Kraken [47] and BLAST analyses indicated that the DNase treatment successfully degraded unpackaged DNA, as only 155 reads (0.14% of the total) with an average length of 1.2kb were derived from the salmon-sperm DNA spike-in (**Fig 4B**). To further reduce the possibility of unpackaged *B. burgdorferi* DNA carryover, we imposed a stringent 5kb read-length cutoff, thus reducing the number of salmon-derived reads to zero and leaving a total of 58,399 reads (**Fig 4C**) with a median length of ∼12.3 kb (**Fig 4D**). Note that we detected a high number of ∼32 kb reads in each replicate which are the approximate size of cp32 prophages (**Fig 4D**, dashed line).

Overall, ∼99.6% of packaged reads >5 kb were classified as *B. burgdorferi* (**Fig 5A**), the majority of which (∼79%) were cp32 isoforms (**Fig 5B**). Cp32-10 and cp32-3 were preferentially packaged (∼32% and ∼25%, respectively) followed by cp32-13 and cp32-5 (each at ∼10%) (**Fig 5B**). Reads mapping to cp32-3, cp32-5, cp32-10, and cp32-13 had a mean coverage of over 1,000× (**Fig 5C**). Cp32-1 reads accounted for only about one percent of all packaged reads (**Fig 5B**) and had lower mean coverage of approximately 36× (**Fig 5C**), suggesting that cp32-1 was not undergoing lytic replication. Read length distributions across cp32s indicate that full-length ∼32 kb molecules were often recovered for cp32-3, cp32-5, and cp32-13, but less frequently for cp32-1 and cp32-10 (**Fig 5D**).

**Figure 5.**
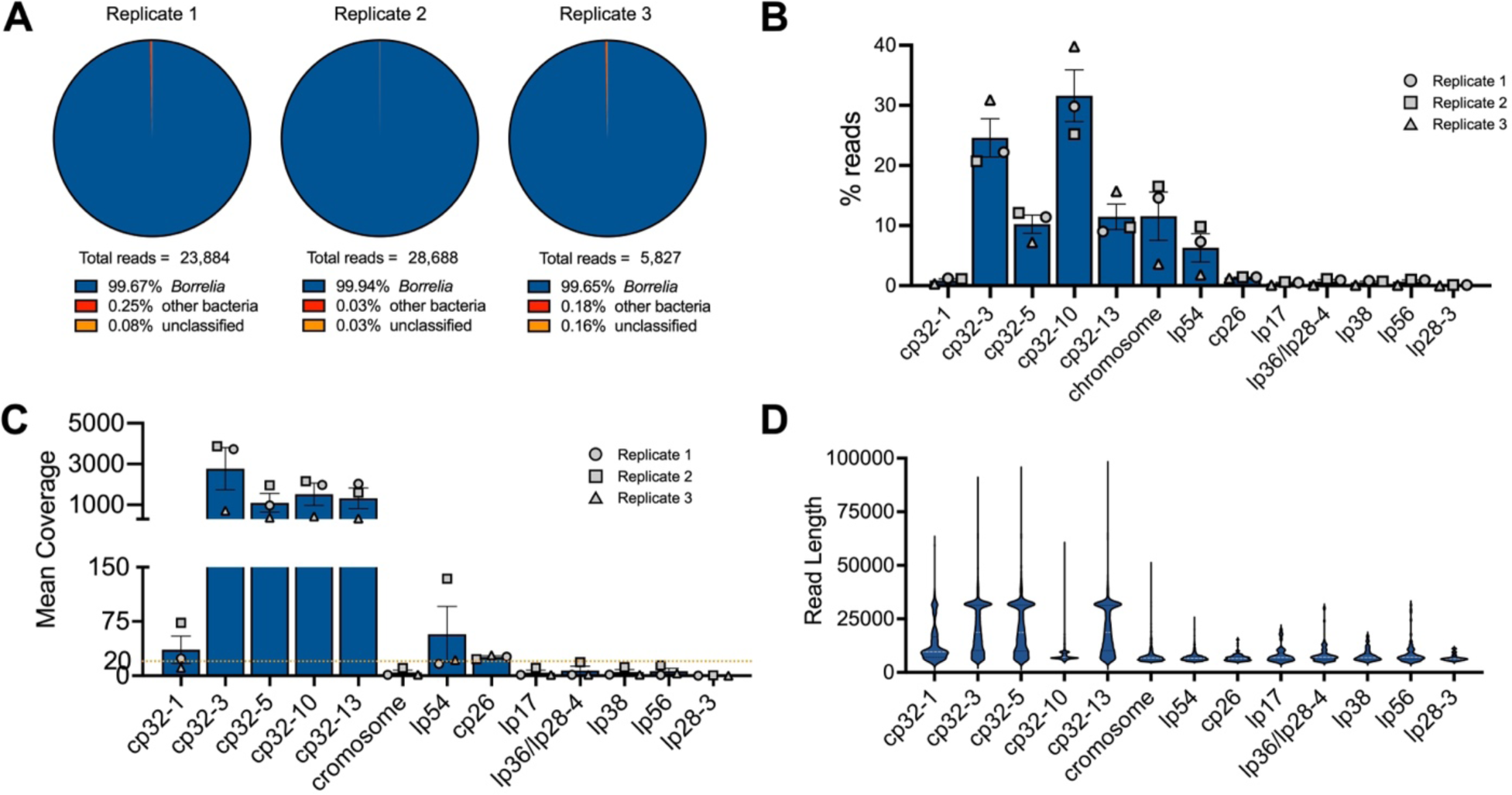
ɸBB-1 virions package cp32 isoforms, chromosome fragments, lp54, and other plasmid DNA. **(A)** Kraken and BLAST were used to determine the taxonomic affiliation of reads >5kb. Note that no eukaryotic reads were identified. **(B and C)** The (B) percent and (C) mean coverage for reads affiliated with the indicated *B. burgdorferi* plasmid or linear chromosome are shown for each replicate. Error bars represent the SE of the mean. **(D)** Read length distributions for the indicated plasmids or chromosome are shown.

Additionally, 11.6% of reads > 5 kb mapped to the linear chromosome and ∼6.3% of reads >5 kb mapped to lp54 (**Fig 5B**). The remaining reads mapped to all the defined genetic elements of *B. burgdorferi* CA-11.2A including plasmids cp26, lp17, lp36/lp28-4, lp38, lp56, and lp28-3 at 1–2% each (**Fig 5B**). *De novo* assembly of packaged reads produced full-length contigs of all cp32s, lp17, cp26, lp36, lp38, lp54, and lp56 (**Fig S1**), suggesting that full-length versions of these plasmids are packaged by ϕBB-1.

Of note, the CA-11.2A genome was reported to contain a unique plasmid, lp36/lp28-4, that is thought to have arisen from the fusion of lp36 with lp28-4 [48]. *De novo* assembly of packaged reads resolved lp36/lp28-4 into individual lp36 and lp28-4 contigs (**Fig S1E and F**). Additionally, whole genome sequencing of our CA-11.2A strain confirmed that lp36 and lp28-4 are separate as no reads that span the lp36-lp28-4 junction were observed and coverage depth was notably different between lp36 and lp28-4 (∼200× vs. 25×, respectively, **Fig S2A**). Furthermore, PCR confirmed the sequencing results (**Fig S2B-D**). These data indicate that the lp36/lp28-4 plasmid is two distinct episomes in our CA-11.2A strain.

Collectively, these results indicate that in addition to cp32 molecules, ϕBB-1 is capable of packaging non-cp32 portions of the *B. burgdorferi* genome. We discuss the major packaged DNA species in the following sections.

**Figure S1.**
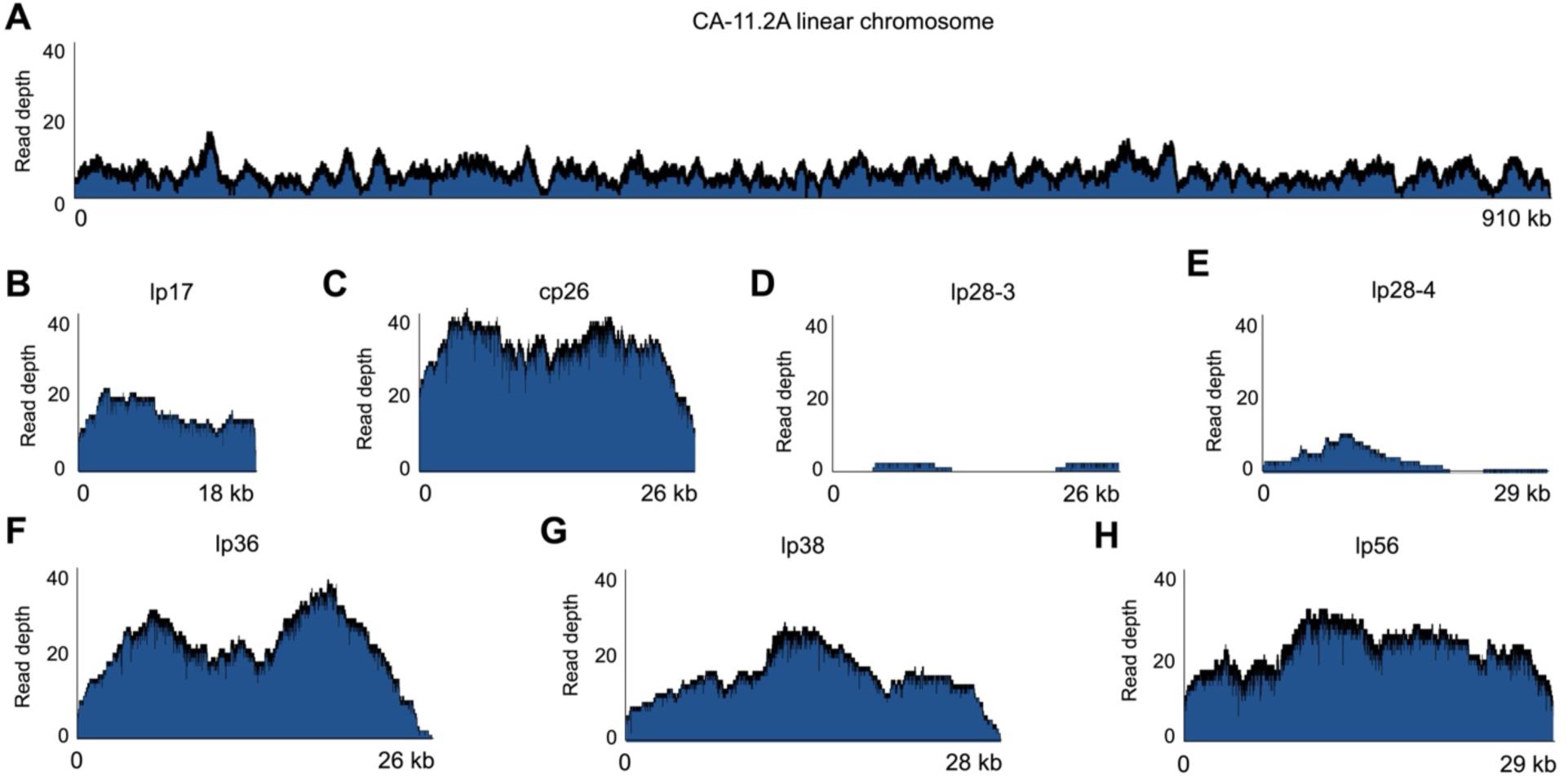
Packaged read depth across the *de novo* CA-11.2A genome assembly. *De novo* assembly of packaged reads >5kb produced the indicated contigs. Read coverage was then mapped to each contig. **(A–H)** Read coverage across the CA-11.2A chromosome or indicated plasmids are shown. Coverage maps for the cp32s and lp54 are shown in Figures 6 and 9, respectively.

**Figure S2.**
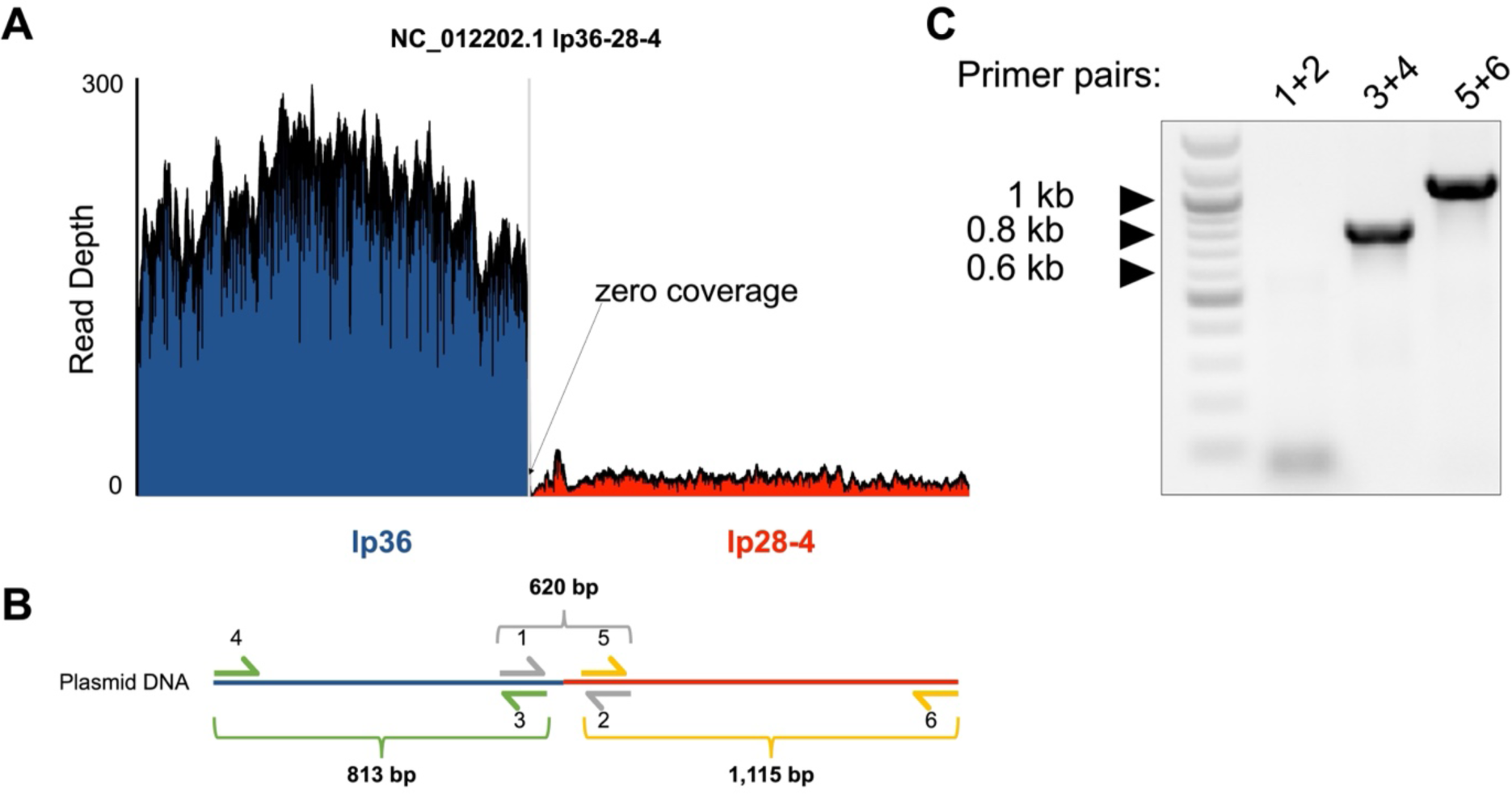
Whole genome sequencing of the CA-11.2A genome reveals that plasmid lp36/lp28-4 resolves into two separate episomes. **(A)** The CA-11.2A genome was sequenced using long-read technology. Reads were aligned to the lp36/lp28-4 reference sequence (NC_012202.1) and read depth plotted. **(B)** Schematic of PCR design. Primers 1 and 2 flank the lp36/lp28-4 junction, with primer 1 annealing to lp36 and primer 2 annealing to lp28-4, creating a 620 bp product if joined. Primers 3 and 4 anneal to lp36 DNA, creating an 813 bp product if present. Primers 5 and 6 anneal to lp28-4 DNA, creating a 1,115-bp product if present. **(C)** The presence or absence of lp36, lp28-4, or lp36/lp28-4 was confirmed by PCR.

### cp32 molecules are linearized near the *erp* locus and packaged via a headful mechanism

Our sequencing data provide insight into how ϕBB-1 packages cp32 molecules. Many phage species package linear double-stranded DNA genomes that circularize after being injected into a host [49]. Because DNA isolated from ϕBB-1 virions is thought to be linearized [33], we used PhageTerm [50] to predict the linear ends of packaged DNA. Native DNA termini are present once per linear DNA molecule, but non-native DNA ends produced during sequencing are distributed randomly along DNA molecules. Thus, reads that start at native DNA terminal positions occur more frequently than anywhere else in the genome. PhageTerm takes advantage of this to resolve DNA termini and predict phage packaging mechanisms [50]. PhageTerm identified the termini of packaged cp32 molecules at approximately 26 kb in a region lying immediately upstream of the *erp* loci (**Fig 6A**). In agreement with the PhageTerm results, when packaged reads were used to map the physical ends of packaged cp32 molecules, a sharp boundary in coverage depth is observed upstream of the *erp* loci in all cp32s (**Fig 6B–F**). Notably, the intergenic region upstream of the *erp* loci is conserved across the cp32 isoforms found in diverse strains of Lyme disease spirochetes (**Fig 6G**) [15] and the linear cp32 ends identified by long-read sequencing converge at the same conserved terminal sequence motif (**Fig 6H**).

**Fig 6.**
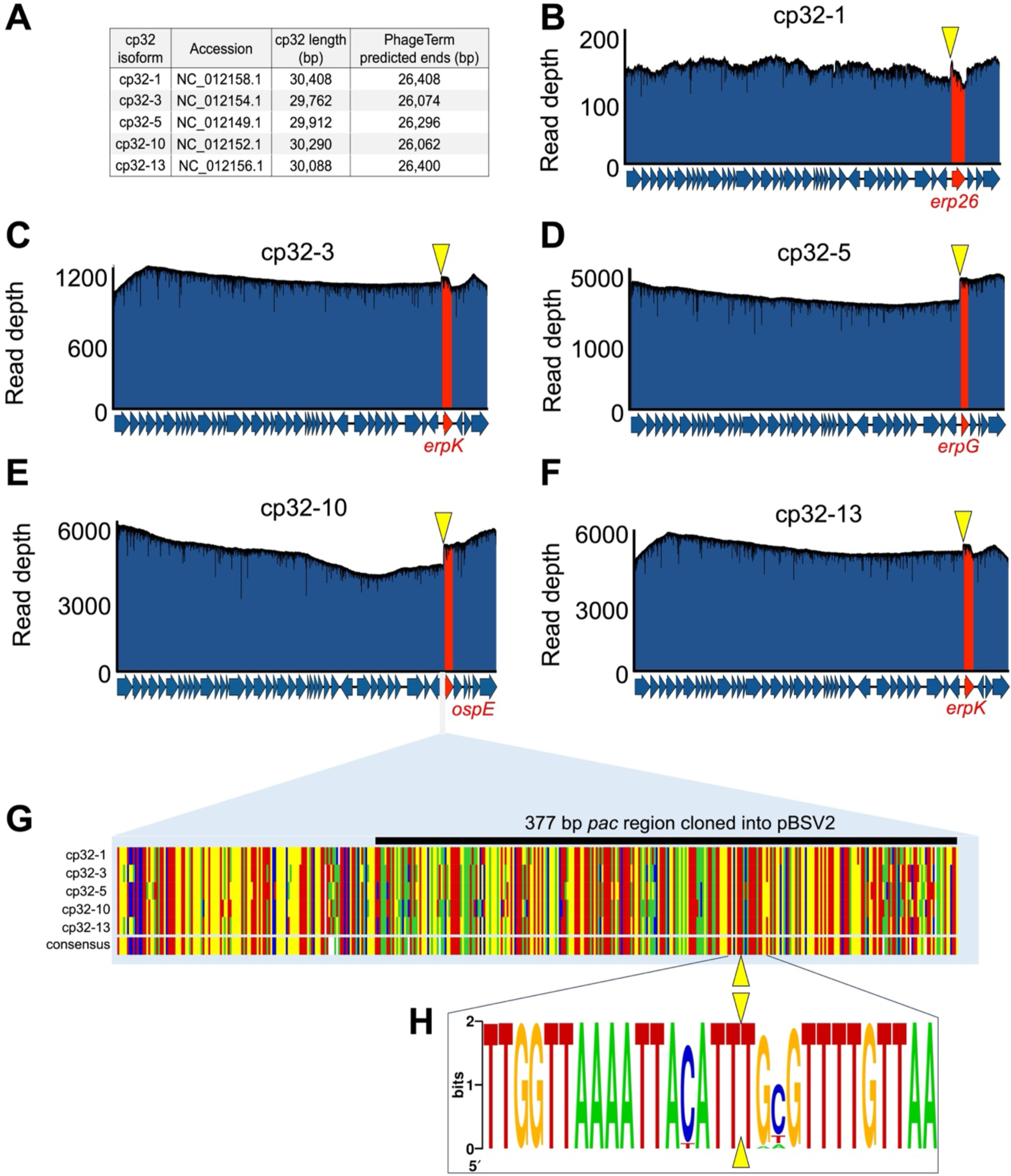
cp32s are linearized upstream of the *erp* loci. **(A)** PhageTerm was used to predict the linear ends of packaged cp32 molecules. **(B–F)** Nanopore reads were mapped to the indicated cp32s. Note the sharp boundary just upstream of the *erp* loci (highlighted in red). The yellow triangles indicate the PhageTerm predicted linear ends. **(G)** Alignments of the intergenic region upstream of the *erp* loci is shown for each cp32. Colors indicating A, T, C, or G are shown in panel H. The black line indicates the *pac* region that was cloned into a shuttle vector, as described in Figure 7. **(H)** A nucleic acid logo was constructed from 207 cp32 sequence alignments. Yellow triangles indicate the linear end of cp32 isoforms as predicted by PhageTerm and confirmed by long-read sequencing.

PhageTerm predicts that cp32s are packaged by a headful mechanism which supports the previously proposed headful genome packaging mechanism for cp32s [42]. Phages that use the headful packaging mechanism generate a concatemer containing several head-to-tail copies of their genome (**Fig 7A**). During headful packaging, a cut is made at a defined packaging site (*pac* site) and a headful (a little more than a full genome) of linear phage DNA is packaged. Once a headful is achieved, the phage genome is cut at non-defined sites, resulting in variable cut positions and size variation in packaged DNA, which we observe in packaged cp32 reads downstream of the initial cut site (**Fig 6B–F**).

**Figure 7.**
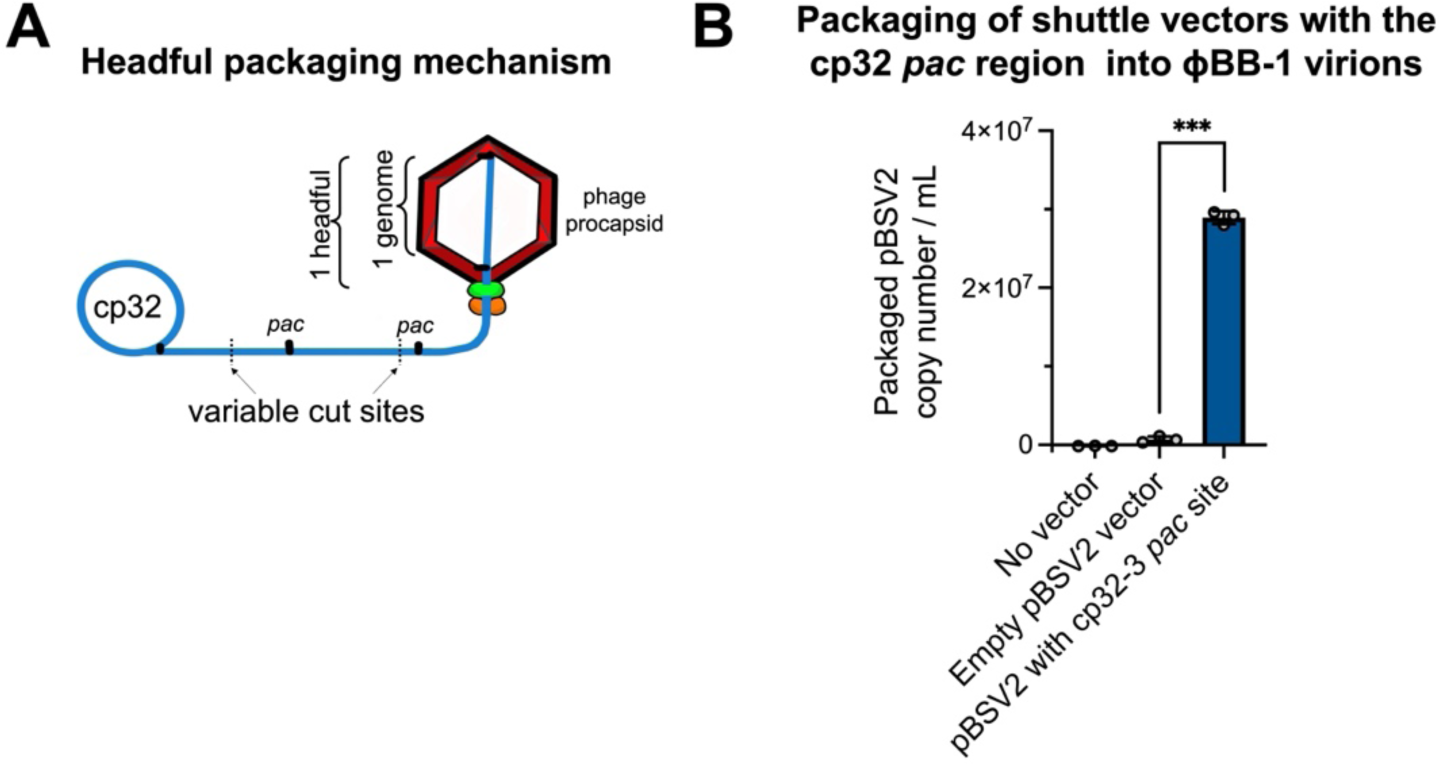
Shuttle vectors containing the cp32 *pac* region are preferentially packaged into ϕBB-1 virions. **(A)** Schematic depicting the headful genome packaging mechanism. **(B)** After ethanol induction, ϕBB-1 virions were collected from CA-11.2A cells not carrying plasmid pBSV2 (No vector), cells transformed with empty pBSV2, or cells transformed with pBSV2 with the cp32-3 *pac* site (see Fig 6G for the cloned *pac* region). Copy numbers of pBSV2 packaged into ϕBB-1 virions were measured by qPCR. Data are the SE of the mean of three experiments, \*\*\**p*<0.001.

Our results suggest that the cp32 *pac* site is upstream of the *erp* loci. If the cp32 *pac* site is in this region, then DNA molecules containing the *pac* sequence are expected to be packaged into ϕBB-1 virions. To test this, we cloned the putative cp32-3 *pac* site (**Fig 6G**, black bar) into a derivative of the pBSV2 shuttle vector that lacks the promoter and MCS [51], transformed *B. burgdorferi* strain CA-11.2A, and induced lytic ϕBB-1 replication with 5% ethanol. Supernatants containing virions were collected, filtered, treated with chloroform, and DNase treated as described above. pBSV2 shuttle vector copy numbers were measured by qPCR using primers that target the pBSV2 kanamycin resistance (*kan*) cassette. To control for possible chromosomal DNA contamination, qPCR was also performed using primers targeting the chromosomal *flaB* gene. Final packaged pBSV2 copy numbers were calculated by subtracting *flaB* copy numbers from pBSV2 (*kan* cassette) copy numbers.

Copy numbers of packaged pBSV2 encoding the cp32-3 *pac* site were significantly (*p*<0.001) higher compared to virions collected from the supernatants of cells carrying an empty pBSV2 vector (**Fig 7B**), indicating that DNA molecules that contain the *pac* site are preferentially packaged by ϕBB-1 virions.

### The cp32 prophages have conserved motifs that occur in a specific arrangement not found in other DNA sequences packaged by ϕBB-1 virions

To identify motif(s) that may be shared between the cp32s and other genomic elements that are packaged into ϕBB-1 virions (*e.g.,* lp54), we first used an iterative BLAST search to identify distantly homologous DNA sequences (Fig 8). A non-redundant list of these diverse DNA sequences were then used as an input dataset for sequence motif discovery via MEME [52]. All five cp32 isoforms found in *B. burgdorferi* CA-11.2A have the same specific arrangement of conserved sequence motifs around the *pac* region (**Fig 8A and B**) and these are conserved in cp32 isoforms across *B. burgdorferi* (**Fig 8C**). However, significant matches to these motifs were not identified in other CA-11.2A genetic elements packaged by ϕBB-1 (**Supplementary Data file 1**), suggesting that packaging of non-cp32 DNA may occur spontaneously or through different mechanisms.

**Figure 8.**
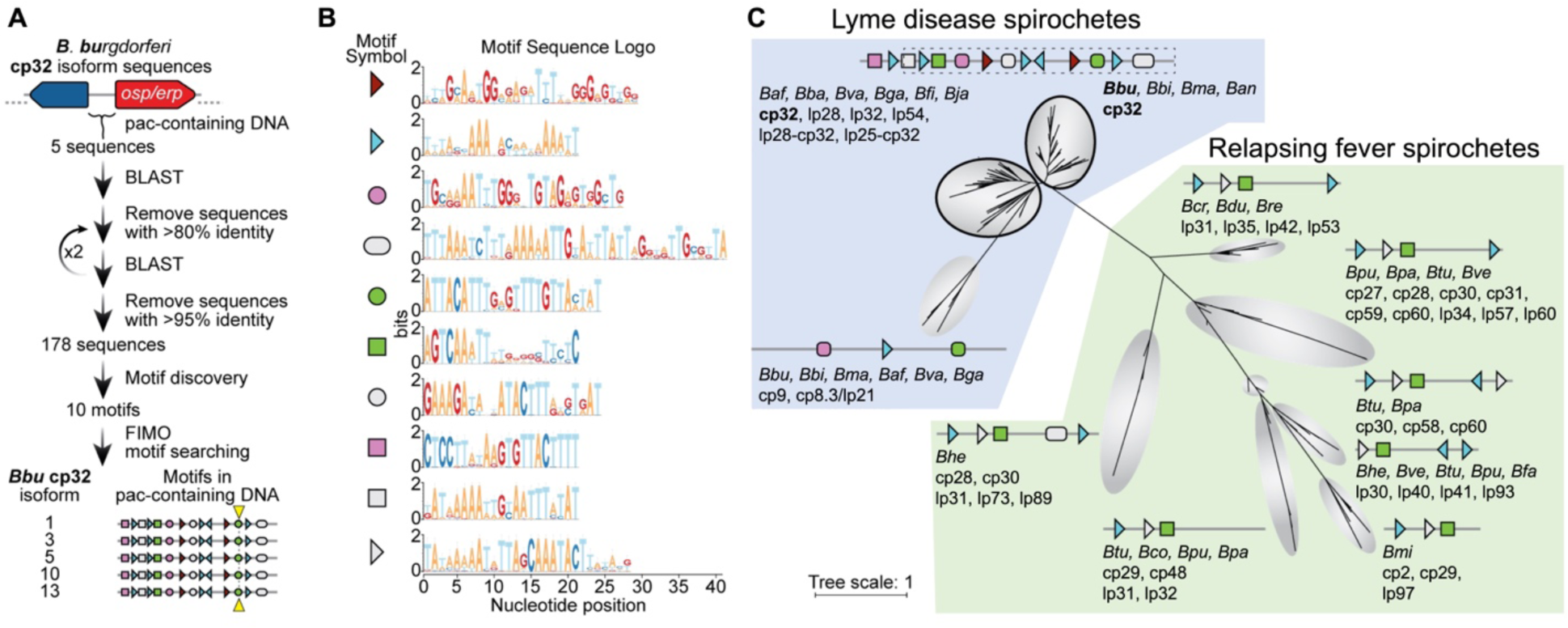
Cp32 prophages have conserved motifs that occur in a specific arrangement around the *pac* site. **(A)** Outline of bioinformatic strategy to identify motifs enriched in the *pac*-containing DNA sequence of cp32 isoforms. All *B. burgdorferi* cp32 isoforms have the same motifs in the *pac* region. The cp32 cut site is indicated by the yellow triangle. **(B)** Sequence logos of the motifs identified in panel A and schematized in panel C. Nine of the top ten motifs occur at least once in the *pac*-containing region of cp32 DNA sequences. Motifs represented with right or left facing triangles often occur as direct and/or indirect repeats. **(C)** Phylogenetic tree of non-redundant DNA sequences with homology to *B. burgdorferi* cp32 *pac*-region identified in panel A. For each clade, the bacterial species and type of plasmid are listed. For clarity in the figure, bacterial species names have been truncated to a three letter abbreviation consisting of the first letter of the genus and the first two letters of the species (*Borrelia afzelii, Baf; Borrelia andersonii, Ban; Borrelia bavariensis, Bba; Borrelia bissettiae, Bbi; Borrelia burgdorferi, Bbu; Borrelia coriaceae, Bco; Borrelia crocidurae, Bcr; Borrelia duttoni, Bdu; Borrelia fainii, Bfa; Borrelia finlandensis, Bfi; Borrelia garinii, Bga; Borrelia hermsii, Bhe; Borrelia japonica, Bja; Borrelia mayonii, Bma; Borrelia miyamotoi, Bmi; Borrelia parkeri, Bpa; Borrelia puertoricensis, Bpu; Borrelia recurrentis, Bre; Borrelia turicatae, Btu; Borrelia valaisiana, Bva; Borrelia venezuelensis, Bve*). There is variability in the motif architecture between sequences within a single clade; however, for clarity, a representative motif architecture discovered by MEME is shown [52]. The top two clades of sequences (outlined in black) are dominated by cp32 isoforms and the cp32 motif architecture, therefore a single motif scheme is shown for these two clades. The region of DNA and motifs cloned into the pBSV2 shuttle vector is outlined in dashes.

The complete or partial arrangement of motifs found around the *pac* site of *B. burgdorferi* cp32 isoforms is conserved in cp32 plasmids and some linear plasmids originating from other Lyme and relapsing fever *Borrelia* (21 species total) (Fig. 8C). The iterative BLAST search also revealed that a diverse set of circular and linear plasmids in a broader set of *Borrelia* species share some of the motifs found in *B. burgdorferi* cp32 isoforms. In total, linear or circular plasmid sequences from 21 different *Borrelia* species (both Lyme disease and relapsing fever spirochetes) had homology to the *B. burgdorferi* cp32 *pac*-containing DNA sequences (**Fig 8C**).

### Deciphering the structure of linear plasmids packaged by ϕBB-1

After the cp32s, lp54 is a major DNA species packaged by ϕBB-1 (**Fig 5C**). Lp54 is a linear plasmid with covalently closed telomeres that is present in all Lyme disease *Borrelia* with about a third of its encoded genes being paralogues to genes encoded on the cp32s [6, 53]. *De novo* assembly of packaged lp54 reads produces a 67.4 kb contig consisting of full-length lp54 (54,021 bp, NC_012194.1) flanked by sequences containing tail-to-tail (7,310 bp) and head-to-head (6,074 bp) junctions (**Fig 9A**). Read depth for lp54 was >100 for most of the contig; however, read depth drops precipitously at both tail-to-tail and head-to-head junctions (**Fig 9A**), suggesting that the telomeres of lp54 interfere with sequencing.

**Figure 9.**
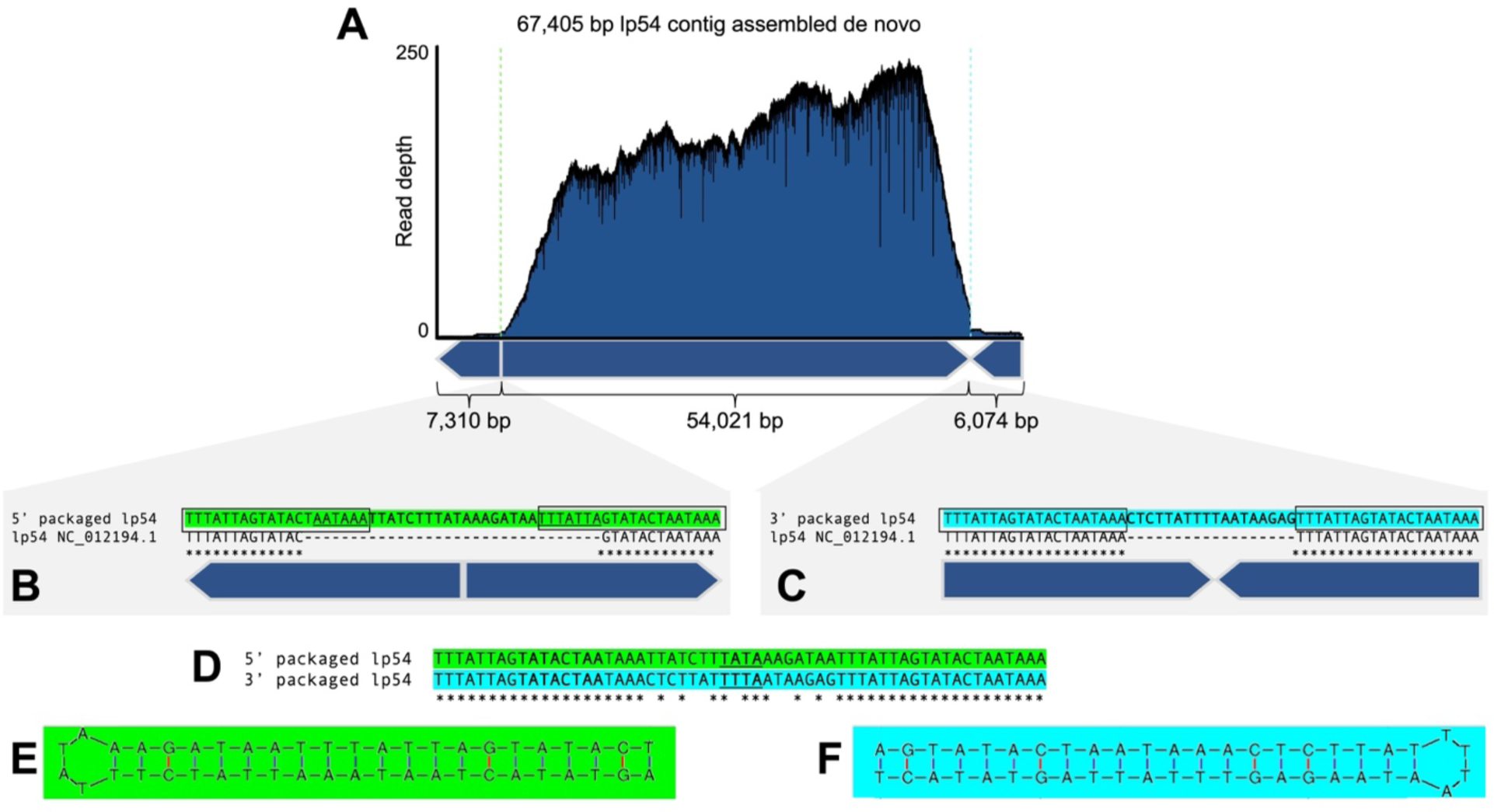
Full-length lp54 with fully resolved telomeres are recovered from ϕBB-1-packaged DNA. **(A)** *De novo* assembly of packaged reads produced a 67,405-bp contig with tail-to-tail and head-to-head junctions. **(B and C)** Sequences at the packaged 5′ junction (green) or the 3′ junction (cyan) are compared to the lp54 reference sequence NC_012194.1. The conserved inverted repeat sequence 5′– TTTATTAGTATACTAATAAA is outlined. **(D)** Alignments of the tail-to-tail and head-to-head junctions reveals a variable 18-bp sequence in between the conserved inverted repeats. **(E and F)** Predicted hairpin structures are shown for each end of lp54. The loop sequence for each hairpin is underlined in panel D.

*B. burgdorferi* telomeres contain inverted repeat sequences [54] and we identified the CA-11.2A lp54 inverted repeat sequence as 5′–TTTATTAGTATACTAATAAA (**Fig 9B and C**, boxed sequences). Our sequencing of the telomeric ends of lp54 extends the reference sequence at the left telomeric end by seven nucleotides (**Fig 9B**, underlined). Further, compared to the lp54 reference sequence, the packaged left and right junction-spanning sequences each encode an additional 18 bp of sequence (**Fig 9B and C**). These sequences, although unique at each end (**Fig 9D**), form perfect hairpin structures (**Fig 9E and F**). Overall, these data suggest that lp54 molecules with complete telomere sequences are packaged into virions. However, whether linear lp54 with covalently closed telomeres or lp54 replication intermediates that contain head-to-head and tail-to-tail junctions are packaged is unclear.

The *de novo* assembly approach applied to lp54 was also successful in resolving the telomeric ends of other linear elements of the CA-11.2A genome, including the linear chromosome and plasmids lp17, lp56, and lp38 (Fig 10). Additionally, we were able to resolve left and right telomeres for lp36 (Fig 10), providing yet further evidence that lp36 is not fused to lp28-4.

**Figure 10.**
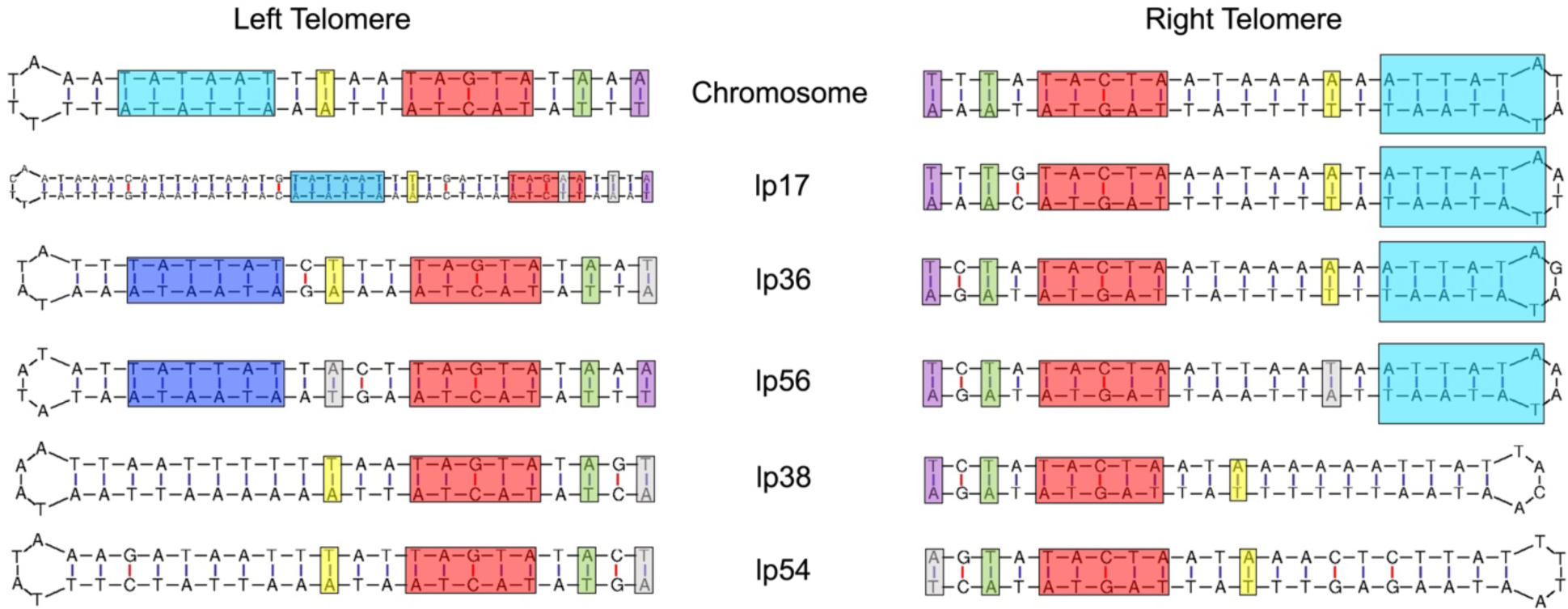
Packaged reads resolve the telomeric ends of the linear chromosome and most linear plasmids in the CA-11.2A genome. Reads spanning tail-to-tail or head-to-head junctions of the linear chromosome or the indicated linear plasmids form perfect hairpin structures. Conserved regulatory elements for each telomere are highlighted [55–60].

## Discussion

In nature, Lyme disease spirochetes exist as diverse populations of closely related bacteria that possess sufficient antigenic variability to allow them to co-infect and reinfect non-naïve vertebrate hosts [61–72]. Moreover, horizontal gene transfer between Lyme disease spirochetes has been extensively documented [19, 73-77]. Nevertheless, the mechanism underlying horizontal genetic exchange among Lyme disease spirochetes has remained undefined. Our study implicates ϕBB-1 in mediating horizontal gene transfer between Lyme disease spirochetes.

Horizontal gene transfer between heterologous spirochetes likely occurs in the tick midgut during and immediately after a blood meal when spirochete replication rates and densities are at their highest. ϕBB-1 replication is also induced in the tick midgut during a bloodmeal [9, 30, 31] with implications for their facilitation of horizontal gene transfer evidenced by homologous recombination between cp32 isoforms [15–17] and the horizontal transfer of cp32s between *Borrelia* strains [21].

Our sequencing data indicate that ϕBB-1 virions package portions of the entire *B. burgdorferi* genome, giving ϕBB-1 the potential to mobilize numerous beneficial alleles during the enzootic cycle via generalized transduction. For example, the circular cp32 prophages are highly conserved across the *Borrelia* genus [26]; however, cp32 isoforms contain variable regions that encode outer membrane lipoproteins such as Bdr, Mlp, and OspE/OspF/Elp, which are known to facilitate the *B. burgdorferi* lifecycle [24, 26, 27, 78]. The linear plasmid lp54 encodes the outer membrane lipoproteins OspA and OspB, which are required for *B. burgdorferi* to colonize the tick midgut [79–81]. The outer membrane lipoprotein OspC, which is required for *B. burgdorferi* to infect a vertebrate host, is encoded by the circular plasmid cp26 [61, 76, 82]. These alleles (and many others) are packaged by ϕBB-1, which is consistent with a role for phage-mediated transduction of genes encoding essential membrane lipoproteins between heterologous spirochetes.

In *B. burgdorferi*, the linear chromosome is highly conserved as are the circular plasmids cp32 and cp26 and the linear plasmids lp17, lp38, lp54, and lp56 are all evolutionarily stable [4-6, 16, 83]. However, other plasmids distributed across the genospecies show considerably more variation, encode mostly (87%) pseudogenes, and are thought to be in a state of evolutionary decay [6]. The packaged plasmids for which we recovered full-length contigs include the cp32s, cp26, lp17, lp38, lp54, and lp56 —the same plasmids that are evolutionarily stable across the genospecies [4-6, 16, 83]. These observations suggest that genes encoded on ϕBB-1-packaged plasmids are under positive selection, possibly due to the continuous transduction between Lyme disease spirochetes during the enzootic cycle.

In addition to providing evidence that ϕBB-1 virions package large portions of the *B. burgdorferi* genome, our study provides insight into ϕBB-1 virion structure and identifies virion proteins present in ϕBB-1. Using mass spectrometry-based proteomics, we confirm that putative capsid and structural genes encoded by the cp32s, such as the major capsid protein P06, are indeed translated and assembled into mature ϕBB-1 virions.

Our long-read sequencing studies indicate that ϕBB-1 packages full-length linear cp32 molecules via a headful mechanism using *pac* sites. The headful packaging mechanism is used by numerous phages and was first described for *E. coli* phage T4 in 1967 [84]. After injecting linear DNA into a new host, the phage genome re-circularizes before continuing its replication cycle. Genes encoded near the ends of linear phage genomes are subject to copy number variation and recombination as the phage genome re-circularizes [85]. Our data suggest that the conversion of linear cp32 molecules into circular cp32 molecules occurs in the vicinity of the *erp* locus, which would facilitate recombination with polymorphic *erp* alleles encoded by other cp32 isoforms in diverse *B. burgdorferi* hosts.

In this study, the packaging of specific cp32 isoforms was biased: cp32-3, cp32-5, cp32-10, and cp32-13 were predominantly packaged while cp32-1 was rarely packaged. This result is consistent with observations by Wachter *et al.* where cp32 isoform copy number and transcriptional activity were not uniform across all cp32 isoforms in *B. burgdorferi* strain B31: cp32-1, cp32-3, and cp32-6 were predominantly induced (highest copy numbers) and had the highest transcriptional activity while cp32-9 was not induced and was transcriptionally inactive [9]. Variability in the *pac* region or other regulatory elements involved in cp32 induction may explain why different cp32 isoforms replicate and/or are packaged at different rates. On the other hand, the motifs that are found most broadly in the *pac* region (*e.g.,* **Fig 8C**, blue triangle and green square) may represent binding sites for conserved host factors that are present in all *Borrelia* species whereas the other motifs may represent protein-binding sites or regulatory sequences that are specific to given prophage or plasmids.

In the intergenic region upstream of the *erp* loci, we identified a 377-bp region that contains the cp32 *pac* signal. Introducing the cp32 *pac* region to a shuttle vector facilitated the packaging of the shuttle vector into ϕBB-1 virions. Our identification of the cp32 *pac* site will be useful for the engineering of recombinant DNA that can be packaged into virions that infect spirochetes, giving ϕBB-1 the potential for use as a tool for the genetic dissection and manipulation of Lyme disease spirochetes.

After the cp32s, lp54 was the most frequently packaged plasmid. This may be related to the evolutionary origins of lp54: about one-third of the genes encoded by lp54 are paralogous to cp32-encoded genes and lp54 is thought to have emerged from an ancient recombination event between a cp32 and a linear plasmid [6]. In addition, lp54 encodes putative phage proteins including a porin (BBA74) [86] and phage capsid proteins that are highly conserved across the genospecies [87], which we detected in purified virions by mass spectrometry. While we observed virions with a distinct elongated capsid morphology, virions with a notably smaller capsid morphology have been observed after induction *in vitro* [9, 32, 33]. These observations raise the possibility that lp54 may be a prophage, although it is not clear if lp54 produces its own capsids, relies on cp32-encoded capsids, or if both lp54 and cp32 capsid proteins assemble to produce chimeric virions.

Our long-read dataset contained reads that spanned head-to-head and tail-to-tail junctions in lp54. These reads allowed us to define the lp54 telomere sequences; however, whether full-length lp54 molecules are packaged or at which stage of the replication cycle lp54 is packaged is unknown. In *B. burgdorferi*, both the linear chromosome and linear plasmids have covalently closed hairpin telomeres and replicate via a telomere resolution mechanism [56, 58, 88, 89]. Examination of a naturally occurring lp54 dimer in *B. valaisiana* isolate VS116 suggests that a circular head-to-head dimer is produced during lp54 replication prior to telomere resolution and replication completion [90]. Linear, covalently closed lp54 molecules may be packaged or lp54 replication intermediates may be packaged.

As obligate vector-borne bacteria, Lyme disease spirochetes live relatively restrictive lifestyles that might be expected to i) limit their exposure to novel gene pools, ii) enhance reductive evolution, and iii) favor the loss of mobile DNA elements. A role for ϕBB-1 in mediating the transduction of beneficial alleles between heterologous spirochetes in local vector and reservoir host populations may explain why cp32 prophages are ubiquitous not only among Lyme disease spirochetes, but also relapsing fever spirochetes.

## Acknowledgments

We are grateful to Patti Rosa for helpful discussions and to the IDeA National Resource for Quantitative Proteomics Center at the University of Arkansas for their assistance with proteomic analyses of phage virions. PRS is supported by NIH grants R21AI151597 and P30GM140963. MK is supported by NIH grant P20GM103474. DRF is supported by NSF GRFP grant 366502. A.S-F. is a M. Jane Williams and Valerie Vargo Presidential Assistant Professor of Biology and is supported by NIH grants K99GM147842 and R00GM147842, and by the Postdoctoral Enrichment Program Award from the Burroughs Wellcome Fund (G-1021106.01). The funders had no role in study design, data collection and analysis, decision to publish, or preparation of the manuscript. The authors declare no conflicts of interest.

## Methods

### ϕBB-1 induction

*Borrelia burgdorferi* B31 or CA-11.2A was grown in BSK-II growth medium to 7 × 10^7^/mL and centrifuged at 6,000 × *g*, 10 min., 35°C to pellet cells, which were resuspended in fresh media to a density of 2 × 10^8^/mL. EtOH was added to a final concentration of 5% and the resuspended culture was incubated at 35°C for an additional 2 hours to induce phage production. The induced culture was then centrifuged at 6,000 × *g*, 10 min, 35°C and the pellet was resuspended in fresh media to a density of 5 × 10^7^/mL after which it was incubated at 35°C for 72 hours to produce phage. After 72 hours, the culture was centrifuged at 6,000 × *g* for 10 min to remove cells and the phage-containing supernatant was filtered twice through 0.2 µm filters before storage at 4°C.

### cp32 qPCR

For qPCR, 100 µL of filtered culture supernatant was mixed with 20ul of chloroform to eliminate remaining intact cells and then centrifuged to separate the phases. 80 µL of the aqueous phase was transferred to a new tube, mixed with 0.8 µL of 100X DNaseI reaction buffer (1M Tris-HCl pH 7.5, 250 mM MgCl_2_, 50 mM CaCl_2_) and DNase treated with 0.8U DNaseI for 1 hour at 37°C. Following DNase treatment, supernatants were mixed with 20 µl chloroform to inactivate DNase, spun to separate phases and the aqueous phase added directly to a qPCR reaction (0.5 µL treated supernatant/10 µL total reaction volume). qPCR was performed using SsoAdvanced Universal Inhibitor-Tolerant SYBR green supermix (BioRad, Hercules, CA) following maufacturer’s instructions, primers that target a conserved cp32 intergenic region between *bbp08* and *bbp09* (5’-CTTTACACATATCAAGACCTTAAC, 5’-CAAACCACCCAATTTCCAATTCC) and the *flaB* gene to control for *B. burgdorferi* chromosomal DNA contamination (5’-TCTTTTCTCTGGTGAGGGAGCT, 5’-TCCTTCCTGTTGAACACCCTCT) [91] at an empirically determined annealing temperature of 55°C. Absolute cp32 and *flaB* copy numbers were calculated from a standard curve generated using a cloned copy of the target sequences. To estimate phage number for CA-11.2A and correct for any remaining unpackaged cp32 plasmids, five times the number of detected *flaB* copies was subtracted from the absolute cp32 starting quantity.

### ϕBB-1 virion purification for DNA extraction

Centrifuged, filtered phage supernatants were treated with 1/10^th^ volume of chloroform to lyse any remaining cells and chloroform was allowed to separate at 4°C overnight. The aqueous layer was transferred to a new vessel and mixed with saturated ammonium sulfate to a final concentration of 50%. NaOH was slowly added during ammonium sulfate addition to maintain pH based on the BSK-II phenol red indicator and the final pH was adjusted to 7.5. Precipitations were incubated overnight 4°C and then centrifuged at 10,000 × g for 30 minutes (4°C) to collect phage pellets. Precipitated phages were gently resuspended in SM buffer overnight at 4°C.

### ϕBB-1 electron microscopy imaging

Purified virions (3–4 μl) were absorbed to the surface of freshly glow-discharged, formvar-coated 200 mesh copper grids and negatively stained with 5 μl of 2% methylamine vanadate (Nanoprobes, Yaphank, NY) prior to viewing on a Hitachi HT7700 transmission electron microscope (Hitachi-High-Technologies Corporation, Tokyo, Japan).

### ϕBB-1 virion proteomics

Purified virions (200 µg total protein) were reduced, alkylated, and purified by chloroform/methanol extraction prior to digestion with sequencing grade modified porcine trypsin (Promega). Peptides were separated on an Acquity BEH C18 column (100 x 1.0 mm, Waters) using an UltiMate 3000 UHPLC system (Thermo). Peptides were eluted by a 50 min gradient from 99:1 to 60:40 buffer A:B ratio (Buffer A = 0.1% formic acid, 0.5% acetonitrile. Buffer B = 0.1% formic acid, 99.9% acetonitrile). Eluted peptides were ionized by electrospray (2.4 kV) followed by mass spectrometric analysis on an Orbitrap Eclipse Tribrid mass spectrometer (Thermo) using multi-notch MS3 parameters. MS data were acquired using the FTMS analyzer over a range of 375 to 1500 m/z. Up to 10 MS/MS precursors were selected for HCD activation with normalized collision energy of 65 kV, followed by acquisition of MS3 reporter ion data using the FTMS analyzer over a range of 100-500 m/z. Proteins were identified and quantified using MaxQuant (Max Planck Institute) TMT MS3 reporter ion quantification with a parent ion tolerance of 2.5 ppm and a fragment ion tolerance of 0.5 Da.

### Packaged ϕBB-1 DNA purification

For DNA extractions, phage were collected and precipitated as described above, with the addition of a DNase treatment prior to ammonium precipitation. The aqueous phage of chloroform supernatants were mixed with 1/100^th^ volume 100X DNAse buffer and 1U/mL DNase I followed by incubated at 37°C for 3 hours and by 4°C overnight. For samples subjected to population sequencing, high molecular-weight salmon sperm DNA (1.7 µg/mL, a concentration that approximates the amount of DNA released by 3 × 10^8^ lysed bacterial cells per milliliter of media) was added prior to DNAse digestion to assess carryover of DNA contained outside of phage capsids.

After ammonium sulfate precipitation and resuspension of phage pellets in SM buffer, EDTA was added to a final concentration of 5 mM and SDS to a final concentration of 0.5%. After addition of 20 µg/mL RNAse and incubation at room temperature for 20 minutes, phage capsids were digested with 200ug/mL proteinase K at 55°C for 1 hour. Samples were extracted twice with an equal volume of phenol-chloroform-isoamyl alcohol (25:24:1) followed by a single extraction with an equal volume of chloroform-isoamyl alcohol (24:1) using Qiagen Maxtract High Density medium (Qiagen, Hilden, Germany). NaCl was added to 300 mM and DNA was precipitated with 2.5 volumes of 100% EtOH at −20°C overnight. DNA was pelleted by centrifugation (14,000 × g for 20 min at 4°C), washed 3X with 70% EtOH and re-spun for 20 min, at 14,000 × g 4°C. The DNA pellet was gently air-dried followed by resuspension in 10mM Tris-HCl, pH 8.5 at 4°C overnight.

### Nanopore sequencing

Sequencing libraries were prepared according to manufacturer’s instructions using library kit SQK-LSK112, native barcoding kit SQK-NBD112.24 and 500 ng of purified phage DNA (Oxoford Nanopore, Oxford, UK). Libraries were sequenced on a MinION MK1-B using a FLO-MIN112 flowcell and default settings until pores were exhausted. Basecalling and demultiplexing was performed with Guppy 6.4.6 using the super high accuracy (SUP) model (dna_r10.4_e8.1_sup.cfg) and default parameters. Run quality control measures were checked with MinIONQC (v1.4.1) [92] and FastQC (v0.11.9). Adaptor trimming was performed using s (v0.2.4) [93]. Reads were deposited in the NCBI BioProject database accession <u>PRJNA1059007</u> and in Supplementary Data File 2.

### Sequence analysis pipeline

Adapter-trimmed long-reads with quality scores ≥7 were used to isolate ≥ 5kb reads using Filtlong (v0.2.1). ≥5kb reads were mapped to the reference *B. burgdorferi* CA-11.2A genome (RefSeq assembly: GCF_000172315.2) with minimap2 (v2.26-r1175) [94]. Primary mapping reads with MAPQ >20 were isolated by contig, filtered, and converted to final file formats using Samtools (v1.17) [95] and SeqKit (v2.5.1) [96]. Read statistics for each replicate were graphed and viewed using GraphPad Prism (v10.1.1). For each contig, *de novo* assemblies were created using Trycycler (v0.5.4) [97], which relied on input assemblies from Flye (v2.9.2-b1786) [98], Raven (v1.8.3) [99], and Minimap2/Miniasm/Minipolish (v2.26-r1175/v0.3-r179/v0.1.2) [94, 100]. The long-read *de novo* assemblies were then polished with short reads using Minipolish (v0.1.2) [100]. The telomeres of the linear chromosome and linear plasmids were manually identified in SnapGene (v5.3.3), and the hairpin structures were predicted by the Mfold webserver (http://www.unafold.org/mfold/applications/dna-folding-form.php) [101]. The terminal ends of the cp32 prophage genomes were predicted using PhageTerm through the Galaxy webserver (https://galaxy.pasteur.fr/) [50], via input of the ≥5kb long-read sequences. Coverage maps of the primary mapping or primary and supplementary mapping reads were created by mapping ≥ 5kb long-reads to the *de novo* assembled CA-11.2A genome or the reference *B. burgdorferi* CA-11.2A genome with Minimap2, converted to final file formats using Samtools, and viewed using R (v4.3.2) and ggplot2 (v3.4.4).

### *Pac* site cloning and qPCR

The putative *pac* region from CA-11.2A genomic DNA was amplified using primers 5’-TAGACATGAGCGGCCGCAAGACAAGCTCCTTATAAGTGTTACT-3’ and 5’-ATAGCTAGATGCGGCCGCTTACTCCGTAACTCTAAAGAATAATGC-3’, purified and digested with NotI and cloned into Not-I-digested pBSV2_2 [51] to create a shuttle vector in which the CA-11.2A *pac* region is maintained but cannot be expressed. Vector sequences were verified using long-read sequencing and transformed into CA-11.2A via electroporation [102]. Clones were PCR-screened for maintenance of resident plasmids as previously described using published primers for *B. burgdorferi* cp32-1, cp26, cp32-3 (which target CA-11.2A cp32-5), cp32-6 (which target CA-11.2A cp32-3), lp28-3, lp17, lp54, lp28-4 [103] and CA-11.2A-specific primers for cp32-3 (5’-TGGGTTGTAGAGTGGCTGTG-3’, 5’-TCACCACTTGCGTAATTCTTGC-3’), cp32-10 (5’-TAGAGCAAAGTCTTGAAAAGACAAC-3’, 5’-CCCACGCTTTGTTGAGACC-3’) and cp32-13 (5’-AATCTGGGCTGTAGAGCAGG-3’, 5’-CTGCTCCTGAGGCTCATCC-3’). Clones transformed with *pac* plasmids or the empty vector were grown in triplicate to late-log phase in BSK-II and used to generate phage as described above. Encapsidated vector was measured directly from DNase-treated culture supernatants as described above using qPCR primers that target the *kan* resistance gene on pBSV2_2 (5’-CACCGGATTCAGTCGTCACT-3’, 5’-GATCCTGGTATCGGTCTGCG-3’, 120 bp product). A cloned copy of the *kan* PCR product was used to generate a standard curve for absolute quantification.

### Identification of conserved motifs in *B. burgdorferi* cp32 isoforms

The roughly 430 nucleotides upstream of the *erp26*, *erpK*, *erpG*, *ospE* and *erpK* genes of the *B. burgdorferi* CA-11.2A cp32 isoforms cp32-1, cp32-3, cp32-5, cp32-10 and cp32-13 respectively were used as queries for a discontinuous MegaBLAST against the NCBI Nucleotide collection database. The results from these first five BLASTs were combined and sequence hits with more than 80% identity were removed with CD-HIT [104]. The resulting representative sequences were used as queries for discontinuous MegaBLAST against the NCBI Nucleotide collection (nt) database, and sequence hits with more than 80% identity were removed with CD-HIT [104]. This process was iterated twice more for a total of three MegaBLAST searches with a representative list of 80% identity query sequences. The sequence hits from the final MegaBLASTs were combined and sequences with more than 95% identity were removed with CD-HIT [104], generating a list of 178 sequences. These 178 sequences were used as an input dataset for the MEME webserver [52], with custom parameters of “Maximum Number of Motifs” set to “10”, and “Motif Site Distribution” set to “Any number of sites per sequence”. MEME identified motifs in 160 of the input sequences. The Position Weight Matrices (PWMs) of the 10 motifs identified by MEME were used as inputs for FIMO [105] to search for significant sequence matches (q-value < 0.001) in the *B. burgdorferi* chromosome and the *B. burgdorferi* cp32-1, cp32-3, cp32-5, cp32-10, cp32-13, cp26, lp17, lp54 plasmid DNA sequences. The cp32 isoforms had nine highly conserved sequence motifs, some motifs present in multiple copies and arranged in a conserved architecture. The cp26, lp17, lp54 and chromosome sequences did not contain this conserved architecture of nine motifs (see Supplementary Data file 1). The sequence logo of each motif was generated by taking the sequence fragments that MEME used to make each PWM, and submitting these sequence fragments to the WebLogo 3.0 webserver [106]. The iterative discontinuous MegaBLAST searches had introduced eukaryotic sequence fragments into the list of 178 non-redundant sequences, suggesting that the search likely reached an endpoint and found most of the related sequences in the NCBI database. To generate a phylogenetic tree, eukaryotic sequence fragments were first removed, and the remaining 149 non-redundant sequences were aligned using the MAFFT webserver [107], with custom parameters of “Direction of nucleotide sequences” set to “Adjust direction according to the first sequence”, and “Strategy” set to E-INS-2. The resulting alignment was used as input for the IQ-TREE webserver [108, 109], with the following command-line: path_to_iqtree -s *.fasta -st DNA -m TEST -bb 1000 -alrt 1000. TreeViewer was used to display the phylogenetic tree [110].

## References

1. Mead PS. Epidemiology of Lyme disease. Infect Dis Clin North Am. 2015;29(2):187–210. Epub 2015/05/23. doi: 10.1016/j.idc.2015.02.010. PubMed PMID: 25999219.

2. Adrion ER, Aucott J, Lemke KW, Weiner JP. Health care costs, utilization and patterns of care following Lyme disease. PLoS One. 2015;10(2):e0116767. Epub 2015/02/05. doi: 10.1371/journal.pone.0116767. PubMed PMID: 25650808; PubMed Central PMCID: PMC4317177.

3. Radolf JD, Strle K, Lemieux JE, Strle F. Lyme Disease in Humans. Curr Issues Mol Biol. 2021;42:333–84. Epub 20201211. doi: 10.21775/cimb.042.333. PubMed PMID: 33303701; PubMed Central PMCID: PMCPMC7946767.

4. Schwartz I, Margos G, Casjens SR, Qiu WG, Eggers CH. Multipartite Genome of Lyme Disease Borrelia: Structure, Variation and Prophages. Curr Issues Mol Biol. 2021;42:409–54. Epub 20201217. doi: 10.21775/cimb.042.409. PubMed PMID: 33328355.

5. Casjens SR, Mongodin EF, Qiu WG, Luft BJ, Schutzer SE, Gilcrease EB, et al. Genome stability of Lyme disease spirochetes: comparative genomics of Borrelia burgdorferi plasmids. PLoS One. 2012;7(3):e33280. Epub 20120314. doi: 10.1371/journal.pone.0033280. PubMed PMID: 22432010; PubMed Central PMCID: PMCPMC3303823.

6. Casjens S, Palmer N, van Vugt R, Huang WM, Stevenson B, Rosa P, et al. A bacterial genome in flux: the twelve linear and nine circular extrachromosomal DNAs in an infectious isolate of the Lyme disease spirochete Borrelia burgdorferi. Mol Microbiol. 2000;35(3):490–516. Epub 2000/02/15. PubMed PMID: 10672174.

7. Samuels DS, Lybecker MC, Yang XF, Ouyang Z, Bourret TJ, Boyle WK, et al. Gene Regulation and Transcriptomics. Curr Issues Mol Biol. 2021;42:223–66. Epub 20201210. doi: 10.21775/cimb.042.223. PubMed PMID: 33300497; PubMed Central PMCID: PMCPMC7946783.

8. Fraser CM, Casjens S, Huang WM, Sutton GG, Clayton R, Lathigra R, et al. Genomic sequence of a Lyme disease spirochaete, Borrelia burgdorferi. Nature. 1997;390(6660):580-6. doi: 10.1038/37551. PubMed PMID: 9403685.

9. Wachter J, Cheff B, Hillman C, Carracoi V, Dorward DW, Martens C, et al. Coupled induction of prophage and virulence factors during tick transmission of the Lyme disease spirochete. Nat Commun. 2023;14(1):198. Epub 20230113. doi: 10.1038/s41467-023-35897-3. PubMed PMID: 36639656; PubMed Central PMCID: PMCPMC9839762.

10. Qiu WG, Martin CL. Evolutionary genomics of Borrelia burgdorferi sensu lato: findings, hypotheses, and the rise of hybrids. Infect Genet Evol. 2014;27:576–93. Epub 20140403. doi: 10.1016/j.meegid.2014.03.025. PubMed PMID: 24704760; PubMed Central PMCID: PMCPMC4299872.

11. Brisson D, Drecktrah D, Eggers CH, Samuels DS. Genetics of Borrelia burgdorferi. Annu Rev Genet. 2012;46:515–36. Epub 2012/09/15. doi: 10.1146/annurev-genet-011112-112140. PubMed PMID: 22974303; PubMed Central PMCID: PMCPMC3856702.

12. Qiu WG, Bosler EM, Campbell JR, Ugine GD, Wang IN, Luft BJ, et al. A population genetic study of Borrelia burgdorferi sensu stricto from eastern Long Island, New York, suggested frequency-dependent selection, gene flow and host adaptation. Hereditas. 1997;127(3):203–16. doi: 10.1111/j.1601-5223.1997.00203.x. PubMed PMID: 9474903.

13. Haven J, Vargas LC, Mongodin EF, Xue V, Hernandez Y, Pagan P, et al. Pervasive recombination and sympatric genome diversification driven by frequency-dependent selection in Borrelia burgdorferi, the Lyme disease bacterium. Genetics. 2011;189(3):951–66. Epub 20110902. doi: 10.1534/genetics.111.130773. PubMed PMID: 21890743; PubMed Central PMCID: PMCPMC3213364.

14. Combs MA, Tufts DM, Adams B, Lin YP, Kolokotronis SO, Diuk-Wasser MA. Host adaptation drives genetic diversity in a vector-borne disease system. PNAS Nexus. 2023;2(8):pgad234. Epub 20230808. doi: 10.1093/pnasnexus/pgad234. PubMed PMID: 37559749; PubMed Central PMCID: PMCPMC10408703.

15. Brisson D, Zhou W, Jutras BL, Casjens S, Stevenson B. Distribution of cp32 prophages among Lyme disease-causing spirochetes and natural diversity of their lipoprotein-encoding erp loci. Appl Environ Microbiol. 2013;79(13):4115–28. Epub 20130426. doi: 10.1128/AEM.00817-13. PubMed PMID: 23624478; PubMed Central PMCID: PMCPMC3697573.

16. Casjens SR, Gilcrease EB, Vujadinovic M, Mongodin EF, Luft BJ, Schutzer SE, et al. Plasmid diversity and phylogenetic consistency in the Lyme disease agent Borrelia burgdorferi. BMC Genomics. 2017;18(1):165. Epub 20170215. doi: 10.1186/s12864-017-3553-5. PubMed PMID: 28201991; PubMed Central PMCID: PMCPMC5310021.

17. Margos G, Hepner S, Mang C, Marosevic D, Reynolds SE, Krebs S, et al. Lost in plasmids: next generation sequencing and the complex genome of the tick-borne pathogen Borrelia burgdorferi. BMC Genomics. 2017;18(1):422. Epub 2017/06/01. doi: 10.1186/s12864-017-3804-5. PubMed PMID: 28558786; PubMed Central PMCID: PMCPMC5450258.

18. Barbour AG, Travinsky B. Evolution and distribution of the ospC Gene, a transferable serotype determinant of Borrelia burgdorferi. MBio. 2010;1(4). Epub 20100928. doi: 10.1128/mBio.00153-10. PubMed PMID: 20877579; PubMed Central PMCID: PMCPMC2945197.

19. Marconi RT, Samuels DS, Landry RK, Garon CF. Analysis of the distribution and molecular heterogeneity of the ospD gene among the Lyme disease spirochetes: evidence for lateral gene exchange. J Bacteriol. 1994;176(15):4572–82. doi: 10.1128/jb.176.15.4572-4582.1994. PubMed PMID: 7913928; PubMed Central PMCID: PMCPMC196277.

20. Stevenson B, Casjens S, Rosa P. Evidence of past recombination events among the genes encoding the Erp antigens of Borrelia burgdorferi. Microbiology (Reading). 1998;144 ( Pt 7):1869–79. doi: 10.1099/00221287-144-7-1869. PubMed PMID: 9695920.

21. Stevenson B, Miller JC. Intra- and interbacterial genetic exchange of Lyme disease spirochete erp genes generates sequence identity amidst diversity. J Mol Evol. 2003;57(3):309–24. Epub 2003/11/25. doi: 10.1007/s00239-003-2482-x. PubMed PMID: 14629041.

22. Touchon M, Moura de Sousa JA, Rocha EP. Embracing the enemy: the diversification of microbial gene repertoires by phage-mediated horizontal gene transfer. Curr Opin Microbiol. 2017;38:66–73. Epub 20170517. doi: 10.1016/j.mib.2017.04.010. PubMed PMID: 28527384.

23. Brissette CA, Haupt K, Barthel D, Cooley AE, Bowman A, Skerka C, et al. Borrelia burgdorferi infection-associated surface proteins ErpP, ErpA, and ErpC bind human plasminogen. Infect Immun. 2009;77(1):300–6. Epub 2008/11/13. doi: 10.1128/IAI.01133-08. PubMed PMID: 19001079; PubMed Central PMCID: PMCPMC2612283.

24. Brissette CA, Cooley AE, Burns LH, Riley SP, Verma A, Woodman ME, et al. Lyme borreliosis spirochete Erp proteins, their known host ligands, and potential roles in mammalian infection. Int J Med Microbiol. 2008;298 Suppl 1:257–67. Epub 20080111. doi: 10.1016/j.ijmm.2007.09.004. PubMed PMID: 18248770; PubMed Central PMCID: PMCPMC2596196.

25. Stevenson B, Bono JL, Schwan TG, Rosa P. Borrelia burgdorferi erp proteins are immunogenic in mammals infected by tick bite, and their synthesis is inducible in cultured bacteria. Infect Immun. 1998;66(6):2648–54. Epub 1998/05/29. doi: 10.1128/IAI.66.6.2648-2654.1998. PubMed PMID: 9596729; PubMed Central PMCID: PMCPMC108251.

26. Stevenson B, Zuckert WR, Akins DR. Repetition, conservation, and variation: the multiple cp32 plasmids of Borrelia species. J Mol Microbiol Biotechnol. 2000;2(4):411–22. Epub 2000/11/15. PubMed PMID: 11075913.

27. Kenedy MR, Lenhart TR, Akins DR. The role of Borrelia burgdorferi outer surface proteins. FEMS Immunol Med Microbiol. 2012;66(1):1–19. Epub 20120521. doi: 10.1111/j.1574-695X.2012.00980.x. PubMed PMID: 22540535; PubMed Central PMCID: PMCPMC3424381.

28. Stevenson B, Casjens S, Rosa P. Evidence of past recombination events among the genes encoding the Erp antigens of Borrelia burgdorferi. Microbiology (Reading). 1998;144 ( Pt 7):1869–79. doi: 10.1099/00221287-144-7-1869. PubMed PMID: 9695920.

29. Qiu WG, Schutzer SE, Bruno JF, Attie O, Xu Y, Dunn JJ, et al. Genetic exchange and plasmid transfers in Borrelia burgdorferi sensu stricto revealed by three-way genome comparisons and multilocus sequence typing. Proc Natl Acad Sci U S A. 2004;101(39):14150–5. Epub 20040916. doi: 10.1073/pnas.0402745101. PubMed PMID: 15375210; PubMed Central PMCID: PMCPMC521097.

30. Sapiro AL, Hayes BM, Volk RF, Zhang JY, Brooks DM, Martyn C, et al. Longitudinal map of transcriptome changes in the Lyme pathogen Borrelia burgdorferi during tick-borne transmission. Elife. 2023;12. Epub 20230714. doi: 10.7554/eLife.86636. PubMed PMID: 37449477.

31. Tokarz R, Anderton JM, Katona LI, Benach JL. Combined effects of blood and temperature shift on Borrelia burgdorferi gene expression as determined by whole genome DNA array. Infect Immun. 2004;72(9):5419–32. doi: 10.1128/IAI.72.9.5419-5432.2004. PubMed PMID: 15322040; PubMed Central PMCID: PMCPMC517457.

32. Eggers CH, Kimmel BJ, Bono JL, Elias AF, Rosa P, Samuels DS. Transduction by phiBB-1, a bacteriophage of Borrelia burgdorferi. J Bacteriol. 2001;183(16):4771–8. doi: 10.1128/JB.183.16.4771-4778.2001. PubMed PMID: 11466280; PubMed Central PMCID: PMCPMC99531.

33. Eggers CH, Samuels DS. Molecular evidence for a new bacteriophage of Borrelia burgdorferi. J Bacteriol. 1999;181(23):7308–13. Epub 1999/11/26. PubMed PMID: 10572135; PubMed Central PMCID: PMCPMC103694.

34. Zhang H, Marconi RT. Demonstration of cotranscription and 1-methyl-3-nitroso-nitroguanidine induction of a 30-gene operon of Borrelia burgdorferi: evidence that the 32-kilobase circular plasmids are prophages. J Bacteriol. 2005;187(23):7985–95. Epub 2005/11/18. doi: 10.1128/JB.187.23.7985-7995.2005. PubMed PMID: 16291672; PubMed Central PMCID: PMCPMC1291276.

35. Fillol-Salom A, Alsaadi A, Sousa JAM, Zhong L, Foster KR, Rocha EPC, et al. Bacteriophages benefit from generalized transduction. PLoS Pathog. 2019;15(7):e1007888. Epub 2019/07/06. doi: 10.1371/journal.ppat.1007888. PubMed PMID: 31276485; PubMed Central PMCID: PMCPMC6636781.

36. Zinder ND, Lederberg J. Genetic exchange in Salmonella. J Bacteriol. 1952;64(5):679–99. doi: 10.1128/jb.64.5.679-699.1952. PubMed PMID: 12999698; PubMed Central PMCID: PMCPMC169409.

37. Budzik JM, Rosche WA, Rietsch A, O’Toole GA. Isolation and characterization of a generalized transducing phage for Pseudomonas aeruginosa strains PAO1 and PA14. J Bacteriol. 2004;186(10):3270–3. Epub 2004/05/06. doi: 10.1128/JB.186.10.3270-3273.2004. PubMed PMID: 15126493; PubMed Central PMCID: PMCPMC400619.

38. Mandecki W, Krajewska-Grynkiewicz K, Klopotowski T. A quantitative model for nonrandom generalized transduction, applied to the phage P22-Salmonella typhimurium system. Genetics. 1986;114(2):633–57. Epub 1986/10/01. PubMed PMID: 3021574; PubMed Central PMCID: PMCPMC1202961.

39. Ebel-Tsipis J, Botstein D, Fox MS. Generalized transduction by phage P22 in Salmonella typhimurium. I. Molecular origin of transducing DNA. J Mol Biol. 1972;71(2):433–48. Epub 1972/11/14. doi: 10.1016/0022-2836(72)90361-0. PubMed PMID: 4564486.

40. Chiang YN, Penades JR, Chen J. Genetic transduction by phages and chromosomal islands: The new and noncanonical. PLoS Pathog. 2019;15(8):e1007878. Epub 20190808. doi: 10.1371/journal.ppat.1007878. PubMed PMID: 31393945; PubMed Central PMCID: PMCPMC6687093.

41. Eggers CH, Gray CM, Preisig AM, Glenn DM, Pereira J, Ayers RW, et al. Phage-mediated horizontal gene transfer of both prophage and heterologous DNA by varphiBB-1, a bacteriophage of Borrelia burgdorferi. Pathog Dis. 2016;74(9). Epub 2016/11/05. doi: 10.1093/femspd/ftw107. PubMed PMID: 27811049.

42. Eggers CH. Identification and characterization of a bacteriophage of Borrelia burgdorferi Graduate Student Theses, Dissertations, & Professional Papers 10590. 2000.

43. Rumnieks J, Fuzik T, Tars K. Structure of the Borrelia bacteriophage phiBB1 procapsid. J Mol Biol. 2023:168323. Epub 20231020. doi: 10.1016/j.jmb.2023.168323. PubMed PMID: 37866476.

44. Eggers CH, Casjens S, Hayes SF, Garon CF, Damman CJ, Oliver DB, et al. Bacteriophages of spirochetes. J Mol Microbiol Biotechnol. 2000;2(4):365-73. Epub 2000/11/15. PubMed PMID: 11075907.

45. Neubert U, Schaller M, Januschke E, Stolz W, Schmieger H. Bacteriophages induced by ciprofloxacin in a Borrelia burgdorferi skin isolate. Zentralbl Bakteriol. 1993;279(3):307–15. doi: 10.1016/s0934-8840(11)80363-4. PubMed PMID: 8219501.

46. Gillis M, De Ley J, De Cleene M. The determination of molecular weight of bacterial genome DNA from renaturation rates. Eur J Biochem. 1970;12(1):143–53. doi: 10.1111/j.1432-1033.1970.tb00831.x. PubMed PMID: 4984994.

47. Wood DE, Salzberg SL. Kraken: ultrafast metagenomic sequence classification using exact alignments. Genome Biol. 2014;15(3):R46. Epub 20140303. doi: 10.1186/gb-2014-15-3-r46. PubMed PMID: 24580807; PubMed Central PMCID: PMCPMC4053813.

48. Schutzer SE, Fraser-Liggett CM, Casjens SR, Qiu WG, Dunn JJ, Mongodin EF, et al. Whole-genome sequences of thirteen isolates of Borrelia burgdorferi. J Bacteriol. 2011;193(4):1018–20. Epub 20101008. doi: 10.1128/JB.01158-10. PubMed PMID: 20935092; PubMed Central PMCID: PMCPMC3028687.

49. Casjens SR, Gilcrease EB. Determining DNA packaging strategy by analysis of the termini of the chromosomes in tailed-bacteriophage virions. Methods Mol Biol. 2009;502:91–111. doi: 10.1007/978-1-60327-565-1_7. PubMed PMID: 19082553; PubMed Central PMCID: PMCPMC3082370.

50. Garneau JR, Depardieu F, Fortier LC, Bikard D, Monot M. PhageTerm: a tool for fast and accurate determination of phage termini and packaging mechanism using next-generation sequencing data. Sci Rep. 2017;7(1):8292. Epub 20170815. doi: 10.1038/s41598-017-07910-5. PubMed PMID: 28811656; PubMed Central PMCID: PMCPMC5557969.

51. Takacs CN, Kloos ZA, Scott M, Rosa PA, Jacobs-Wagner C. Fluorescent Proteins, Promoters, and Selectable Markers for Applications in the Lyme Disease Spirochete Borrelia burgdorferi. Appl Environ Microbiol. 2018;84(24). Epub 2018/10/14. doi: 10.1128/AEM.01824-18. PubMed PMID: 30315081; PubMed Central PMCID: PMCPMC6275353.

52. Bailey TL, Boden M, Buske FA, Frith M, Grant CE, Clementi L, et al. MEME SUITE: tools for motif discovery and searching. Nucleic Acids Res. 2009;37(Web Server issue):W202–8. Epub 20090520. doi: 10.1093/nar/gkp335. PubMed PMID: 19458158; PubMed Central PMCID: PMCPMC2703892.

53. Casjens S. Evolution of the linear DNA replicons of the Borrelia spirochetes. Curr Opin Microbiol. 1999;2(5):529–34. doi: 10.1016/s1369-5274(99)00012-0. PubMed PMID: 10508719.

54. Tourand Y, Bankhead T, Wilson SL, Putteet-Driver AD, Barbour AG, Byram R, et al. Differential telomere processing by Borrelia telomere resolvases in vitro but not in vivo. J Bacteriol. 2006;188(21):7378–86. Epub 20060825. doi: 10.1128/JB.00760-06. PubMed PMID: 16936037; PubMed Central PMCID: PMCPMC1636258.

55. Tourand Y, Kobryn K, Chaconas G. Sequence-specific recognition but position-dependent cleavage of two distinct telomeres by the Borrelia burgdorferi telomere resolvase, ResT. Mol Microbiol. 2003;48(4):901–11. doi: 10.1046/j.1365-2958.2003.03485.x. PubMed PMID: 12753185.

56. Kobryn K, Chaconas G. Hairpin Telomere Resolvases. Microbiol Spectr. 2014;2(6). doi: 10.1128/microbiolspec.MDNA3-0023-2014. PubMed PMID: 26104454.

57. Huang WM, Robertson M, Aron J, Casjens S. Telomere exchange between linear replicons of Borrelia burgdorferi. J Bacteriol. 2004;186(13):4134–41. doi: 10.1128/JB.186.13.4134-4141.2004. PubMed PMID: 15205414; PubMed Central PMCID: PMCPMC421586.

58. Chaconas G, Stewart PE, Tilly K, Bono JL, Rosa P. Telomere resolution in the Lyme disease spirochete. Embo J. 2001;20(12):3229–37. doi: 10.1093/emboj/20.12.3229. PubMed PMID: 11406599; PubMed Central PMCID: PMCPMC150187.

59. Casjens S, Murphy M, DeLange M, Sampson L, van Vugt R, Huang WM. Telomeres of the linear chromosomes of Lyme disease spirochaetes: nucleotide sequence and possible exchange with linear plasmid telomeres. Mol Microbiol. 1997;26(3):581–96. doi: 10.1046/j.1365-2958.1997.6051963.x. PubMed PMID: 9402027.

60. Tourand Y, Deneke J, Moriarty TJ, Chaconas G. Characterization and in vitro reaction properties of 19 unique hairpin telomeres from the linear plasmids of the lyme disease spirochete. J Biol Chem. 2009;284(11):7264–72. Epub 20090102. doi: 10.1074/jbc.M808918200. PubMed PMID: 19122193; PubMed Central PMCID: PMCPMC2652300.

61. Wilske B, Preac-Mursic V, Jauris S, Hofmann A, Pradel I, Soutschek E, et al. Immunological and molecular polymorphisms of OspC, an immunodominant major outer surface protein of Borrelia burgdorferi. Infect Immun. 1993;61(5):2182–91. doi: 10.1128/iai.61.5.2182-2191.1993. PubMed PMID: 8478108; PubMed Central PMCID: PMCPMC280819.

62. Probert WS, Crawford M, Cadiz RB, LeFebvre RB. Immunization with outer surface protein (Osp) A, but not OspC, provides cross-protection of mice challenged with North American isolates of Borrelia burgdorferi. J Infect Dis. 1997;175(2):400–5. doi: 10.1093/infdis/175.2.400. PubMed PMID: 9203661.

63. Barthold SW. Specificity of infection-induced immunity among Borrelia burgdorferi sensu lato species. Infect Immun. 1999;67(1):36–42. doi: 10.1128/IAI.67.1.36-42.1999. PubMed PMID: 9864193; PubMed Central PMCID: PMCPMC96274.

64. Brisson D, Vandermause MF, Meece JK, Reed KD, Dykhuizen DE. Evolution of northeastern and midwestern Borrelia burgdorferi, United States. Emerg Infect Dis. 2010;16(6):911–7. doi: 10.3201/eid1606.090329. PubMed PMID: 20507740; PubMed Central PMCID: PMCPMC3086229.

65. Nadelman RB, Hanincova K, Mukherjee P, Liveris D, Nowakowski J, McKenna D, et al. Differentiation of reinfection from relapse in recurrent Lyme disease. N Engl J Med. 2012;367(20):1883–90. doi: 10.1056/NEJMoa1114362. PubMed PMID: 23150958; PubMed Central PMCID: PMCPMC3526003.

66. Bhatia B, Hillman C, Carracoi V, Cheff BN, Tilly K, Rosa PA. Infection history of the blood-meal host dictates pathogenic potential of the Lyme disease spirochete within the feeding tick vector. PLoS Pathog. 2018;14(4):e1006959. Epub 20180405. doi: 10.1371/journal.ppat.1006959. PubMed PMID: 29621350; PubMed Central PMCID: PMCPMC5886588.

67. Kurtti TJ, Munderloh UG, Hughes CA, Engstrom SM, Johnson RC. Resistance to tick-borne spirochete challenge induced by Borrelia burgdorferi strains that differ in expression of outer surface proteins. Infect Immun. 1996;64(10):4148–53. doi: 10.1128/iai.64.10.4148-4153.1996. PubMed PMID: 8926082; PubMed Central PMCID: PMCPMC174350.

68. Piesman J, Dolan MC, Happ CM, Luft BJ, Rooney SE, Mather TN, et al. Duration of immunity to reinfection with tick-transmitted Borrelia burgdorferi in naturally infected mice. Infect Immun. 1997;65(10):4043–7. doi: 10.1128/iai.65.10.4043-4047.1997. PubMed PMID: 9317005; PubMed Central PMCID: PMCPMC175581.

69. Hofmeister EK, Glass GE, Childs JE, Persing DH. Population dynamics of a naturally occurring heterogeneous mixture of Borrelia burgdorferi clones. Infect Immun. 1999;67(11):5709–16. doi: 10.1128/IAI.67.11.5709-5716.1999. PubMed PMID: 10531219; PubMed Central PMCID: PMCPMC96945.

70. Shang ES, Wu XY, Lovett MA, Miller JN, Blanco DR. Homologous and heterologous Borrelia burgdorferi challenge of infection-derived immune rabbits using host-adapted organisms. Infect Immun. 2001;69(1):593–8. doi: 10.1128/IAI.69.1.593-598.2001. PubMed PMID: 11119560; PubMed Central PMCID: PMCPMC97926.

71. Derdakova M, Dudioak V, Brei B, Brownstein JS, Schwartz I, Fish D. Interaction and transmission of two Borrelia burgdorferi sensu stricto strains in a tick-rodent maintenance system. Appl Environ Microbiol. 2004;70(11):6783–8. doi: 10.1128/AEM.70.11.6783-6788.2004. PubMed PMID: 15528545; PubMed Central PMCID: PMCPMC525125.

72. Khatchikian CE, Nadelman RB, Nowakowski J, Schwartz I, Wormser GP, Brisson D. Evidence for strain-specific immunity in patients treated for early lyme disease. Infect Immun. 2014;82(4):1408–13. Epub 20140113. doi: 10.1128/IAI.01451-13. PubMed PMID: 24421042; PubMed Central PMCID: PMCPMC3993397.

73. Dykhuizen DE, Polin DS, Dunn JJ, Wilske B, Preac-Mursic V, Dattwyler RJ, et al. Borrelia burgdorferi is clonal: implications for taxonomy and vaccine development. Proc Natl Acad Sci U S A. 1993;90(21):10163–7. doi: 10.1073/pnas.90.21.10163. PubMed PMID: 8234271; PubMed Central PMCID: PMCPMC47734.

74. Stevenson B, Barthold SW. Expression and sequence of outer surface protein C among North American isolates of Borrelia burgdorferi. FEMS Microbiol Lett. 1994;124(3):367–72. doi: 10.1111/j.1574-6968.1994.tb07310.x. PubMed PMID: 7851744.

75. Jauris-Heipke S, Liegl G, Preac-Mursic V, Rossler D, Schwab E, Soutschek E, et al. Molecular analysis of genes encoding outer surface protein C (OspC) of Borrelia burgdorferi sensu lato: relationship to ospA genotype and evidence of lateral gene exchange of ospC. J Clin Microbiol. 1995;33(7):1860–6. doi: 10.1128/jcm.33.7.1860-1866.1995. PubMed PMID: 7665660; PubMed Central PMCID: PMCPMC228286.

76. Livey I, Gibbs CP, Schuster R, Dorner F. Evidence for lateral transfer and recombination in OspC variation in Lyme disease Borrelia. Mol Microbiol. 1995;18(2):257–69. doi: 10.1111/j.1365-2958.1995.mmi_18020257.x. PubMed PMID: 8709845.

77. Will G, Jauris-Heipke S, Schwab E, Busch U, Rossler D, Soutschek E, et al. Sequence analysis of ospA genes shows homogeneity within Borrelia burgdorferi sensu stricto and Borrelia afzelii strains but reveals major subgroups within the Borrelia garinii species. Med Microbiol Immunol. 1995;184(2):73–80. doi: 10.1007/BF00221390. PubMed PMID: 7500914.

78. Hefty PS, Brooks CS, Jett AM, White GL, Wikel SK, Kennedy RC, et al. OspE-related, OspF-related, and Elp lipoproteins are immunogenic in baboons experimentally infected with Borrelia burgdorferi and in human lyme disease patients. J Clin Microbiol. 2002;40(11):4256–65. Epub 2002/11/01. PubMed PMID: 12409407; PubMed Central PMCID: PMCPMC139709.

79. Tilly K, Bestor A, Rosa PA. Functional Equivalence of OspA and OspB, but Not OspC, in Tick Colonization by Borrelia burgdorferi. Infect Immun. 2016;84(5):1565–73. Epub 20160422. doi: 10.1128/IAI.00063-16. PubMed PMID: 26953324; PubMed Central PMCID: PMCPMC4862709.

80. Battisti JM, Bono JL, Rosa PA, Schrumpf ME, Schwan TG, Policastro PF. Outer surface protein A protects Lyme disease spirochetes from acquired host immunity in the tick vector. Infect Immun. 2008;76(11):5228–37. Epub 20080908. doi: 10.1128/IAI.00410-08. PubMed PMID: 18779341; PubMed Central PMCID: PMCPMC2573341.

81. Yang XF, Pal U, Alani SM, Fikrig E, Norgard MV. Essential role for OspA/B in the life cycle of the Lyme disease spirochete. J Exp Med. 2004;199(5):641–8. Epub 20040223. doi: 10.1084/jem.20031960. PubMed PMID: 14981112; PubMed Central PMCID: PMCPMC2213294.

82. Wang IN, Dykhuizen DE, Qiu W, Dunn JJ, Bosler EM, Luft BJ. Genetic diversity of ospC in a local population of Borrelia burgdorferi sensu stricto. Genetics. 1999;151(1):15–30. doi: 10.1093/genetics/151.1.15. PubMed PMID: 9872945; PubMed Central PMCID: PMCPMC1460459.

83. Lemieux JE, Huang W, Hill N, Cerar T, Freimark L, Hernandez S, et al. Whole genome sequencing of human Borrelia burgdorferi isolates reveals linked blocks of accessory genome elements located on plasmids and associated with human dissemination. PLoS Pathog. 2023;19(8):e1011243. Epub 20230831. doi: 10.1371/journal.ppat.1011243. PubMed PMID: 37651316; PubMed Central PMCID: PMCPMC10470944.

84. Streisinger G, Emrich J, Stahl MM. Chromosome structure in phage t4, iii. Terminal redundancy and length determination. Proc Natl Acad Sci U S A. 1967;57(2):292–5. doi: 10.1073/pnas.57.2.292. PubMed PMID: 16591467; PubMed Central PMCID: PMCPMC335503.

85. Beaulaurier J, Luo E, Eppley JM, Uyl PD, Dai X, Burger A, et al. Assembly-free single-molecule sequencing recovers complete virus genomes from natural microbial communities. Genome Res. 2020;30(3):437–46. Epub 2020/02/23. doi: 10.1101/gr.251686.119. PubMed PMID: 32075851; PubMed Central PMCID: PMCPMC7111524.

86. Skare JT, Champion CI, Mirzabekov TA, Shang ES, Blanco DR, Erdjument-Bromage H, et al. Porin activity of the native and recombinant outer membrane protein Oms28 of Borrelia burgdorferi. J Bacteriol. 1996;178(16):4909–18. doi: 10.1128/jb.178.16.4909-4918.1996. PubMed PMID: 8759855; PubMed Central PMCID: PMCPMC178274.

87. Kasumba IN, Tilly K, Lin T, Norris SJ, Rosa PA. Strict Conservation yet Non-Essential Nature of Plasmid Gene bba40 in the Lyme Disease Spirochete Borrelia burgdorferi. Microbiol Spectr. 2023;11(3):e0047723. Epub 20230403. doi: 10.1128/spectrum.00477-23. PubMed PMID: 37010416; PubMed Central PMCID: PMCPMC10269632.

88. Barbour AG, Garon CF. Linear plasmids of the bacterium Borrelia burgdorferi have covalently closed ends. Science. 1987;237(4813):409-11. doi: 10.1126/science.3603026. PubMed PMID: 3603026.

89. Hinnebusch J, Tilly K. Linear plasmids and chromosomes in bacteria. Mol Microbiol. 1993;10(5):917–22. doi: 10.1111/j.1365-2958.1993.tb00963.x. PubMed PMID: 7934868.

90. Marconi RT, Casjens S, Munderloh UG, Samuels DS. Analysis of linear plasmid dimers in Borrelia burgdorferi sensu lato isolates: implications concerning the potential mechanism of linear plasmid replication. J Bacteriol. 1996;178(11):3357–61. doi: 10.1128/jb.178.11.3357-3361.1996. PubMed PMID: 8655522; PubMed Central PMCID: PMCPMC178094.

91. Pahl A, Kuhlbrandt U, Brune K, Rollinghoff M, Gessner A. Quantitative detection of Borrelia burgdorferi by real-time PCR. J Clin Microbiol. 1999;37(6):1958–63. doi: 10.1128/JCM.37.6.1958-1963.1999. PubMed PMID: 10325354; PubMed Central PMCID: PMCPMC84995.

92. Lanfear R, Schalamun M, Kainer D, Wang W, Schwessinger B. MinIONQC: fast and simple quality control for MinION sequencing data. Bioinformatics. 2019;35(3):523–5. doi: 10.1093/bioinformatics/bty654. PubMed PMID: 30052755; PubMed Central PMCID: PMCPMC6361240.

93. Wick RR, Judd LM, Gorrie CL, Holt KE. Completing bacterial genome assemblies with multiplex MinION sequencing. Microb Genom. 2017;3(10):e000132. Epub 20170914. doi: 10.1099/mgen.0.000132. PubMed PMID: 29177090; PubMed Central PMCID: PMCPMC5695209.

94. Li H. Minimap2: pairwise alignment for nucleotide sequences. Bioinformatics. 2018;34(18):3094–100. doi: 10.1093/bioinformatics/bty191.

95. Danecek P, Bonfield JK, Liddle J, Marshall J, Ohan V, Pollard MO, et al. Twelve years of SAMtools and BCFtools. GigaScience. 2021;10(2). doi: 10.1093/gigascience/giab008.

96. Shen W, Le S, Li Y, Hu F. SeqKit: A Cross-Platform and Ultrafast Toolkit for FASTA/Q File Manipulation. PLOS ONE. 2016;11(10):e0163962. doi: 10.1371/journal.pone.0163962.

97. Wick RR, Judd LM, Cerdeira LT, Hawkey J, Méric G, Vezina B, et al. Trycycler: consensus long-read assemblies for bacterial genomes. Genome Biology. 2021;22(1). doi: 10.1186/s13059-021-02483-z.

98. Kolmogorov M, Yuan J, Lin Y, Pevzner PA. Assembly of long, error-prone reads using repeat graphs. Nature Biotechnology. 2019;37(5):540–6. doi: 10.1038/s41587-019-0072-8.

99. Vaser R, Šikić M. Time- and memory-efficient genome assembly with Raven. Nature Computational Science. 2021;1(5):332–6. doi: 10.1038/s43588-021-00073-4.

100. Wick RR, Holt KE. Benchmarking of long-read assemblers for prokaryote whole genome sequencing. F1000Research. 2021;8:2138. doi: 10.12688/f1000research.21782.4.

101. Zuker M. Mfold web server for nucleic acid folding and hybridization prediction. Nucleic Acids Research. 2003;31(13):3406–15. doi: 10.1093/nar/gkg595.

102. Samuels DS. Electrotransformation of the spirochete Borrelia burgdorferi. Methods Mol Biol. 1995;47:253–9. doi: 10.1385/0-89603-310-4:253. PubMed PMID: 7550741; PubMed Central PMCID: PMCPMC5815860.

103. Bunikis I, Kutschan-Bunikis S, Bonde M, Bergstrom S. Multiplex PCR as a tool for validating plasmid content of Borrelia burgdorferi. J Microbiol Methods. 2011;86(2):243–7. Epub 20110512. doi: 10.1016/j.mimet.2011.05.004. PubMed PMID: 21605603.

104. Fu L, Niu B, Zhu Z, Wu S, Li W. CD-HIT: accelerated for clustering the next-generation sequencing data. Bioinformatics. 2012;28(23):3150–2. Epub 20121011. doi: 10.1093/bioinformatics/bts565. PubMed PMID: 23060610; PubMed Central PMCID: PMCPMC3516142.

105. Grant CE, Bailey TL, Noble WS. FIMO: scanning for occurrences of a given motif. Bioinformatics. 2011;27(7):1017–8. Epub 20110216. doi: 10.1093/bioinformatics/btr064. PubMed PMID: 21330290; PubMed Central PMCID: PMCPMC3065696.

106. Crooks GE, Hon G, Chandonia JM, Brenner SE. WebLogo: a sequence logo generator. Genome Res. 2004;14(6):1188–90. Epub 2004/06/03. doi: 10.1101/gr.849004. PubMed PMID: 15173120; PubMed Central PMCID: PMCPMC419797.

107. Katoh K, Standley DM. MAFFT multiple sequence alignment software version 7: improvements in performance and usability. Mol Biol Evol. 2013;30(4):772–80. Epub 20130116. doi: 10.1093/molbev/mst010. PubMed PMID: 23329690; PubMed Central PMCID: PMCPMC3603318.

108. Nguyen LT, Schmidt HA, von Haeseler A, Minh BQ. IQ-TREE: a fast and effective stochastic algorithm for estimating maximum-likelihood phylogenies. Mol Biol Evol. 2015;32(1):268–74. Epub 20141103. doi: 10.1093/molbev/msu300. PubMed PMID: 25371430; PubMed Central PMCID: PMCPMC4271533.

109. Minh BQ, Nguyen MA, von Haeseler A. Ultrafast approximation for phylogenetic bootstrap. Mol Biol Evol. 2013;30(5):1188–95. Epub 20130215. doi: 10.1093/molbev/mst024. PubMed PMID: 23418397; PubMed Central PMCID: PMCPMC3670741.

110. Bianchini G, Sánchez-Baracaldo P. TreeViewer - Cross-platform software to draw phylogenetic trees. 2023. doi: 10.5281/zenodo.7768343.

